# Chronic hyperactivation of midbrain dopamine neurons causes preferential dopamine neuron degeneration

**DOI:** 10.1101/2024.04.05.588321

**Authors:** Katerina Rademacher, Zak Doric, Dominik Haddad, Aphroditi Mamaligas, Szu-Chi Liao, Rose B. Creed, Kohei Kano, Zac Chatterton, YuHong Fu, Joseph H. Garcia, Victoria Vance, Yoshitaka Sei, Anatol Kreitzer, Glenda M Halliday, Alexandra B. Nelson, Elyssa B. Margolis, Ken Nakamura

## Abstract

Parkinson’s disease (PD) is characterized by the death of substantia nigra (SNc) dopamine (DA) neurons, but the pathophysiological mechanisms that precede and drive their death remain unknown. The activity of DA neurons is likely altered in PD, but we understand little about if or how chronic changes in activity may contribute to degeneration. To address this question, we developed a chemogenetic (DREADD) mouse model to chronically increase DA neuron activity, and confirmed this increase using *ex vivo* electrophysiology. Chronic hyperactivation of DA neurons resulted in prolonged increases in locomotor activity during the light cycle and decreases during the dark cycle, consistent with chronic changes in DA release and circadian disturbances. We also observed early, preferential degeneration of SNc projections, recapitulating the PD hallmarks of selective vulnerability of SNc axons and the comparative resilience of ventral tegmental area axons. This was followed by eventual loss of midbrain DA neurons. Continuous DREADD activation resulted in a sustained increase in baseline calcium levels, supporting a role for increased calcium in the neurodegeneration process. Finally, spatial transcriptomics from DREADD mice examining midbrain DA neurons and striatal targets, and cross-validation with human patient samples, provided insights into potential mechanisms of hyperactivity-induced toxicity and PD. Our results thus reveal the preferential vulnerability of SNc DA neurons to increased neural activity, and support a potential role for increased neural activity in driving degeneration in PD.

## Introduction

In Parkinson’s disease (PD), the loss of substantia nigra pars compacta (SNc) dopamine (DA) neurons leads to severe disruption of circuit dynamics in the basal ganglia. Compensation for DA loss involves changes in the activity of both surviving SNc neurons, and of other downstream neurons in the circuit. Indeed, following lesions of the nigrostriatal pathway in rats, surviving SNc DA neurons are hyperactive (*1*), release additional DA (*2–5*), and have reduced DA reuptake (*2*). Massive loss of DA neurons (*1, 6, 7*), complete loss of mitochondrial complex I activity (*8*), and loss of the mitochondrial PD protein *PINK1* (*9*) can also result in increased burst firing (*10, 11*). Therefore, DA neurons are predisposed to altered activity in the setting of extensive loss or stress, which may drive ongoing disease processes. Some lines of evidence support the potential role of hyperactivity in disease initiation, including increased activity in a subset of DA neurons before their degeneration in MitoPark mice (*12*), increased spontaneous firing in PD patient-derived iPSC dopamine neurons (*13*), and increased activity of nigrostriatal dopamine neurons in genetic models of PD (*9, 14*). Moreover, critical PD disease proteins including α-synuclein, LRRK2, PINK1, and Parkin can influence the level of neural activity (*14–19*). In particular, the normal function of α-synuclein is believed to be regulating neural activity (*20*), further supporting the idea that changes in neural activity may contribute to disease pathophysiology. Circuit level changes may also contribute to adverse DA neuron activity. For instance, evidence from primate models suggests that the subthalamic nucleus, which sends a glutamatergic projection to the SNc, is hyperactive in PD (*21*). While system-level changes may be compensatory and partially restore DA levels and overall motor function, they may also have adverse consequences.

Healthy SNc DA neurons are believed to have immense energetic requirements due to their pacemaking activity, active calcium pumping, unmyelinated or poorly myelinated fibers (*22, 23*), and large axonal arbors (*24*). This large energetic requirement likely accounts for their intrinsic vulnerability to mitochondrial insults, including complex I disruption (*8, 25, 26*) and impairments in mitochondrial dynamics (*27*) and turnover (*28*). Neighboring ventral tegmental area (VTA) neurons are relatively spared in PD (*29, 30*), and this may be due to lower reliance on calcium oscillations for pacemaking (*31*), their ability to buffer calcium more effectively than SNc neurons (*29*), and smaller axon arbors (*32, 33*). Thus, combined with disease-related stress, the metabolic impact of even mild hyperactivity may trigger or accelerate degeneration in SNc DA neurons. In support of this hypothesis, inhibiting the excitatory input from the STN protects SNc DA neurons from 6-OHDA and MPTP toxicity (*34, 35*). However, empirical evidence linking chronic changes in neural activity to the degeneration of SNc DA neurons in PD is lacking. Recordings from putative SNc neurons in PD patients displayed twofold higher burst firing when compared to recordings from healthy rodents and nonhuman primates despite similar mean firing rates, though this data is difficult to interpret without human controls (*10, 36*).

Additionally, changes in calcium dynamics during hyperactivity can also drive metabolic stress. SNc DA neurons rely on voltage-gated Ca_v_1.3 calcium channels to support pacemaking, and blocking these channels is protective against the toxicity of 6-OHDA and MPTP (*37, 38*). While classic excitotoxicity involves cytosolic calcium overload and acute cell death, chronic synaptic hyperactivity results in sublethal stress to mitochondria, promoting calcium dysregulation and dendritic atrophy (*39*). Mitochondrial calcium overload has been observed in the setting of LRRK2 mutation and PINK1 deficiency (*39*). Therefore, altered activity, metabolic stress and calcium overload may all contribute to DA neuron death.

To understand if chronic hyperactivation of DA neurons is sufficient to cause neurodegeneration, we developed a chemogenetic mouse model. Our results indicate that chronically increasing neural activity in midbrain DA neurons results in alteration of circadian locomotion patterns, and prolonged activation leads to selective degeneration of SNc axons and eventual death of midbrain DA neurons. These changes were accompanied by altered intracellular calcium dynamics and transcriptomic changes consistent with calcium dysregulation, supporting a role for increased neural activity in driving neurodegeneration in PD.

## Results

To model a chronic increase in DA neuron activity, as may occur in PD, we used a chemogenetic approach. We first expressed the excitatory DREADD hM3Dq specifically in DA neurons using stereotaxic delivery of Cre-dependent hM3Dq AAV to the SNc and VTA of mice expressing Cre under the dopamine transporter (DAT^IRES^Cre) promoter. Next, we measured the acute behavioral effects of chemogenetic activation of DA neurons. As locomotor output is strongly tied to nigrostriatal DA function (*40–42*), we used home cage wheel running as an *in vivo* proxy for changes in DA function. Mice were single-housed and locomotion was quantified based on wheel rotation. Two weeks after viral injection, we administered clozapine-n-oxide (CNO) by i.p. injection (0.5mg/kg) and confirmed that mice responded with an acute increase in wheel running as an indicator of successful DREADD expression (Figure S1A). The resulting hM3Dq-expressing DAT^IRES^Cre mice were then administered either vehicle (2% sucrose, to compensate for CNO’s bitter flavor (*43*)) or CNO (2% sucrose, 300 mg/L CNO) via drinking water ad libitum for two weeks (Figure 1A). This strategy allowed us to chronically activate SNc and VTA DA neurons.

**Figure 1.**
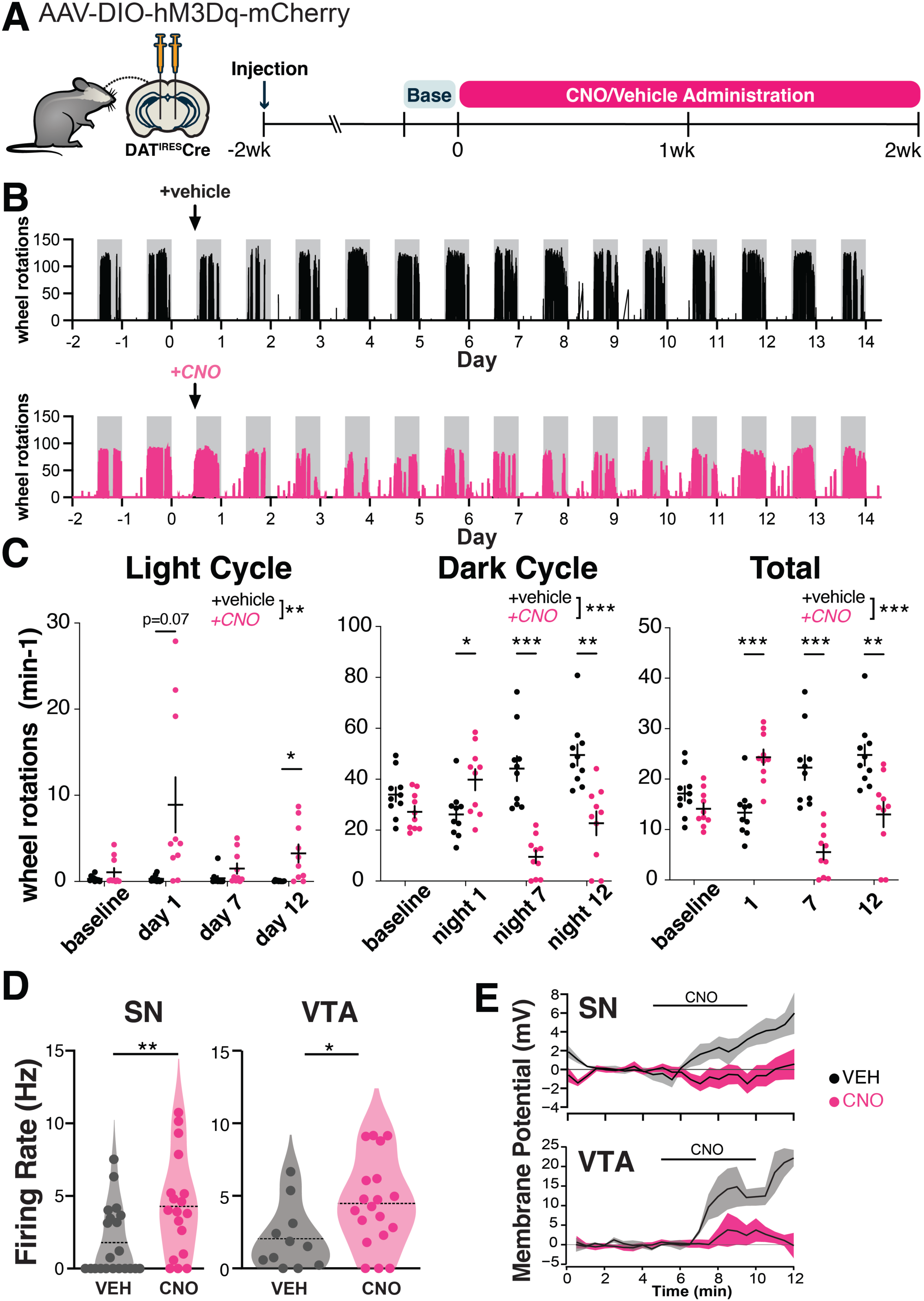
Chronic hM3Dq activation persistently alters the activity of SNc dopamine neurons. **(A)** Graphical illustration summarizing experimental design. Recombinant AAV encoding a conditional allele of the hM3Dq(DREADD)-mCherry was injected bilaterally into the ventral midbrain of 4-5 month-old DAT^IRES^Cre mice. CNO (300 mg/L) or vehicle (2% sucrose in water) was administered *ad libitum* via drinking water for two weeks and the animals were perfused the next day. Changes in locomotion were assessed with running wheels. The two days preceding start of treatment were used as a measure for baseline locomotion. **(B)** Representative traces of wheel usage for animals given control vehicle water (top) or CNO water (bottom). Arrows denote start of treatment. Grey background shading indicates dark cycle hours. **(C)** Mean wheel usage for selected days during the experiment, segregated by light (left), dark (middle), and total (light + dark; right) cycles. n=10 animals/group from 2 independent experiments. *p≤0.05, **p<0.01, ***p<0.001 by two-way ANOVA and Holm-Sidak post hoc test. **(D)** Spontaneous firing rate was measured during the first 2 min of whole cell recordings. *p≤0.05, ** p = 0.01 by t-test or permutation (non-parametric) analysis. **(E)** Time course of responses to bath application of 1 μM CNO *ex vivo* measured in current clamp in neurons from vehicle-treated and CNO-treated mice.

Previous reports indicate that chemogenetic activation of DA neurons leads to increased locomotion (*44*), but the effects of chronic, long-term activation are unknown. During the first day of treatment, CNO-treated animals were more active than controls during the dark cycle, and also strongly trended toward more activity during the light cycle (Figure 1B, 1C). By day 3 of treatment, dark cycle activity markedly decreased in CNO-treated DREADD-expressing mice, and remained decreased for the rest of the treatment time, whereas activity during the light cycle remained increased (Figure 1B, 1C, Figure S1B). Decreased wheel usage in CNO-treated animals may reflect a consequence of circadian disruption. The prolonged changes in wheel activity indicate that the behavioral effects of chemogenetic activation persist throughout the two-week testing period. In contrast, when DAT^IRES^Cre mice were given CNO for four weeks in the absence of the DREADD (CNO alone), there were no significant changes in activity in either light or dark cycles (Figure S1C). These results indicate that CNO alone does not significantly alter mouse activity.

We performed a similar experiment administering CNO for two weeks to DAT^IRES^Cre mice injected with hM3Dq that did not respond to acute CNO IP injection (non-responders; Figure S1D). Non-responders did not show strong initial changes to light or dark cycle wheel usage in the first few days of treatment, but did show persistent decreases in dark cycle usage. Presumably, the lack of early changes reflects either differences in the amount of hM3Dq expressed or the proportion of neurons that are transfected.

To gain insight into the impact of prolonged chemogenetic activation on DA neuron activity, we performed *ex vivo* whole cell recordings in DREADD-expressing SNc neurons in midbrain slices after 7 days of *in vivo* treatment with CNO or vehicle. *In vivo* DREADD activation induced several changes in somatodendritic properties of SNc neurons including a marked decrease in the magnitude of the hyperpolarization activated non-selective cation current *I*_h_ (Figure S2A). It also led to an increase in the spontaneous firing rate (Figure 1D, Figure S2C), indicating a persistent increase in firing rate with CNO treatment. The rate of firing was decreased somewhat in controls relative to historical controls from our lab (*27*), possibly reflecting mild cellular stress from the AAV virus. We did not observe a difference in the coefficient of variation of the interspike interval between treatment groups, thus there was no change in the regularity of pacemaker firing (Figure S2A). *In vivo* CNO led to a more depolarized action potential (AP) peak voltage in spontaneous APs (Figure S2A). CNO treatment did not impact AP threshold voltage or duration (Figure S2A), indicating that AP waveforms were not degraded.

To measure the direct physiological impact of DREADD activation in SNc neurons, we also acutely bath applied 1 uM CNO to these slices. Interestingly, while SNc neurons from CNO-naïve hM3Dq mice depolarized in response to acute CNO, chronic exposure to CNO for one week *in vivo* eliminated this acute response (in neurons from vehicle-treated mice: 4.9 +/− 2.9 mV (n = 5), in neurons from CNO-treated mice: −0.5 +/− 1.0 mV (n = 9); p = 0.05, t test, Figure 1E, Figure S2D). Taken together, these findings indicate that 7d of CNO treatment increased spontaneous firing, but altered hM3Dq function such that acute physiological impacts were no longer apparent. The absence of an acute response may indicate a change in the coupling or availability of the receptors, an adaptation or dysfunction within the neurons, or a homeostatic response to prolonged activation. This may also reflect early stages of toxicity to the CNO-treated DA neurons.

We performed similar recordings in DREADD-expressing VTA neurons. As in the SNc, we found that the spontaneous firing rate was higher with 7-days of *in vivo* DREADD activation (Figure 1D, Figure S2C). Unlike the SNc, the coefficient of variation of the interspike interval was decreased in VTA neurons from the CNO-treated mice, indicating more regular firing patterns (Figure S2B). Also unlike the SNc, VTA neurons did not display changes in *I*_h_ magnitude or changes in AP peak voltage in spontaneous APs following chronic *in vivo* DREADD activation (Figure S2B). *In vivo* treatment did not impact AP threshold voltage, AP peak, AP duration, input resistance, or initial membrane potential in VTA neurons (Figure S2B).

In response to acute bath application of CNO to slices, VTA neurons from CNO-naïve hM3Dq mice depolarized, while those from mice that received chronic 7-day exposure *in vivo* to CNO showed markedly reduced but not fully eliminated responses (in neurons from vehicle-treated mice: 11.2 +/− 8.5 mV (n = 3), in neurons from CNO-treated mice: 2.0 +/− 2.2 mV (n = 7); p = 0.17 t test, Figure 1E, Figure S2D), indicating some retained hM3Dq functionality. Together, these data may reflect resilience to chronic activation in VTA neurons compared to SNc neurons.

### SNc axons are preferentially vulnerable to chronic hM3Dq-activation

To determine if chronic chemogenetic activation of DA neurons for two weeks induces degeneration, we first quantified its effects on axonal integrity in the striatum. Strikingly, compared to vehicle-treated mice, hM3Dq-expressing animals treated with CNO lost ≈40% of their dopaminergic axons in the dorsal striatum, manifest by decreases in both TH immunoreactivity and reporter mCherry immunofluorescence (Figure 2A,B). The same perturbation had a lesser impact on nucleus accumbens and no impact on olfactory tubercle DA afferents. Notably, TH immunoreactivity and mCherry immunofluorescence also decreased in non-responder mice following 2 weeks of treatment (Fig. S3A), although the extent of decrease was somewhat less than in responders. After a shorter 1-week treatment period in a small cohort, we also observed a trend for decreased TH immunoreactivity in the CPu (Figure S3B).

**Figure 2.**
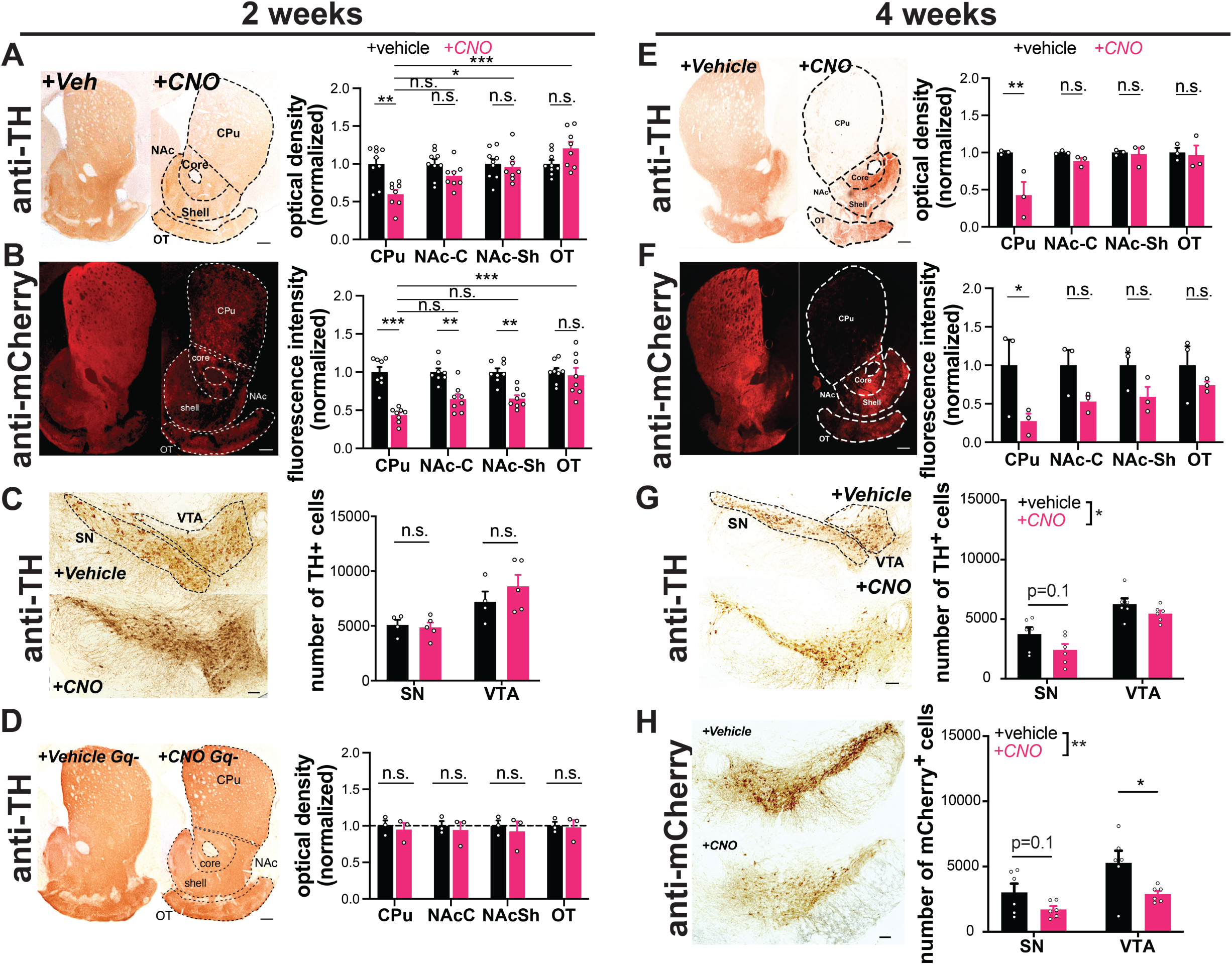
Chronic activation with AAV-hM3Dq-DREADDs is preferentially toxic to nigrostriatal axons. DAT^IRES^Cre mice expressing hM3Dq(DREADD)-mCherry **(A-C, E-H)**, or no virus injection **(D)** in DA neurons. Example images of TH **(A,D,E)** and mCherry **(B,F)** immunoreactivity in striatal sections of mice treated for two or four weeks with vehicle **(left)** or CNO **(right)** via drinking water. DA neuron projection areas in dorsal and ventral striatum are indicated with dotted lines. Quantifications for TH (A,D,E) and mCherry (B,F) optical density at 2 and 4 weeks show preferential loss in CPu. n = 8-9 (A), 8 (B), or 3 (D, E, F) animals/group, 3-5 sections/animal. **(C,G,H)** Images of TH and mCherry (4 weeks only) immunoreactivity in midbrain of vehicle **(top)** or CNO **(bottom)** treated mice at 2 or 4 weeks. SN and VTA regions are indicated with dotted lines. (C,G,H) Stereology estimating the number of TH+ or mCherry+ DA neurons. Chronic CNO treatment of hM3Dq(DREADD)-expressing mice shows a significant decrease in both TH and mCherry immunoreactivity and DA neuron number. n = 4-5 hemispheres (one per mouse, C) or 6 hemispheres (two per mouse, G, H) per group. Scale bars indicate 100μm in the midbrain and 200μm in the striatum. Error bars indicate mean ± SEM. *p<0.05, **p<0.01, ***p < 0.001 by two-way ANOVA and Holm-Sidak *post hoc* test. SN: substantia nigra, VTA: ventral tegmental area, CPu: caudate putamen, NAc-C: nucleus accumbens core, NAc-Sh: nucleus accumbens shell, OT: olfactory tubercule.

We also assessed the impact of two weeks activation on DA neuron survival in the midbrain. With stereological quantification we found no difference in neuron number between CNO-treated animals and vehicle-treated controls in the SNc or VTA (Figure 2C). Therefore, at this time point CNO-treated mice exhibit severe axonal degeneration without DA neuron death.

To ensure that the loss of DA terminals was not due to an off-target effect of CNO, we assessed TH expression in CNO alone animals. These mice did not show a decrease in dopaminergic axons in the caudate putamen (Figure 2D), supporting the conclusion that the axonal degeneration requires hM3Dq activation.

We next assessed the impact of more prolonged activation on degeneration of DA neuron somata. Mice treated with CNO for 4 weeks lost the majority of axons projecting to the caudate putamen (Figure 2E,F), while fibers in the nucleus accumbens and olfactory tubercle were again preferentially spared. Moreover, following 4 weeks of CNO treatment, there was a significant decrease in the overall number of TH+ midbrain (SNc and VTA) DA neurons, assessed by stereology (Figure 2G,H). Although loss of TH reactivity can occur in the absence of neuronal death, there was a similar decrease in the number of mCherry+ DA neurons (labeled by DsRed) in the midbrain. The alignment of these findings supports true loss of DA neurons, although quantitation of mCherry+ DA neurons specifically informs the impact on transduced DA neurons, and this analysis does not account for the variability between injections. Together, these data suggest that, similar to PD progression in humans (*45*), axonal impairment precedes cell body loss in this model. No preferential loss of SNc versus VTA cell bodies was detected at this time point.

### Chronic activation increases baseline population calcium levels in midbrain DA neurons

hM3Dq activation may increase neural activity by increasing intracellular calcium levels (*46*), and increased neural activity should increase intracellular calcium. To determine the impact of 2 weeks of hM3Dq activation on intracellular calcium concentrations in DA neurons, we bred DAT^IRES^Cre mice to GCaMP6f^fl^ mice and injected them bilaterally with the Cre-dependent hM3Dq AAV (Figure 3A). The fluorescent signal was used to guide implantation of an optical fiber in the midbrain of animals (Figure 3B,C), with confirmed hM3Dq expression based on increased cage wheel running in response to i.p. CNO (Figure S1A). Mice were administered CNO or vehicle water and then placed in arenas for 10-min fiber photometry recording sessions every 1-3 days over the 2-week treatment period. Mean fluorescence levels served as a proxy for population level intracellular calcium concentrations. We observed a small trend for increased fluorescent signal the day after starting CNO that persisted for days 3 through 5 but did not reach significance (Figure 3D-F). Interestingly, this was followed by a second, much larger increase in calcium levels between 10-17 days (Figure 3D-F), that occurred in parallel with axon loss (Figure 2A, Figure S3B). To determine if the increase in neural activity was reversible, we performed two additional recording sessions after removing CNO from the drinking water. Although there was a small trend for decreased fluorescence, this did not reach significance.

**Figure 3.**
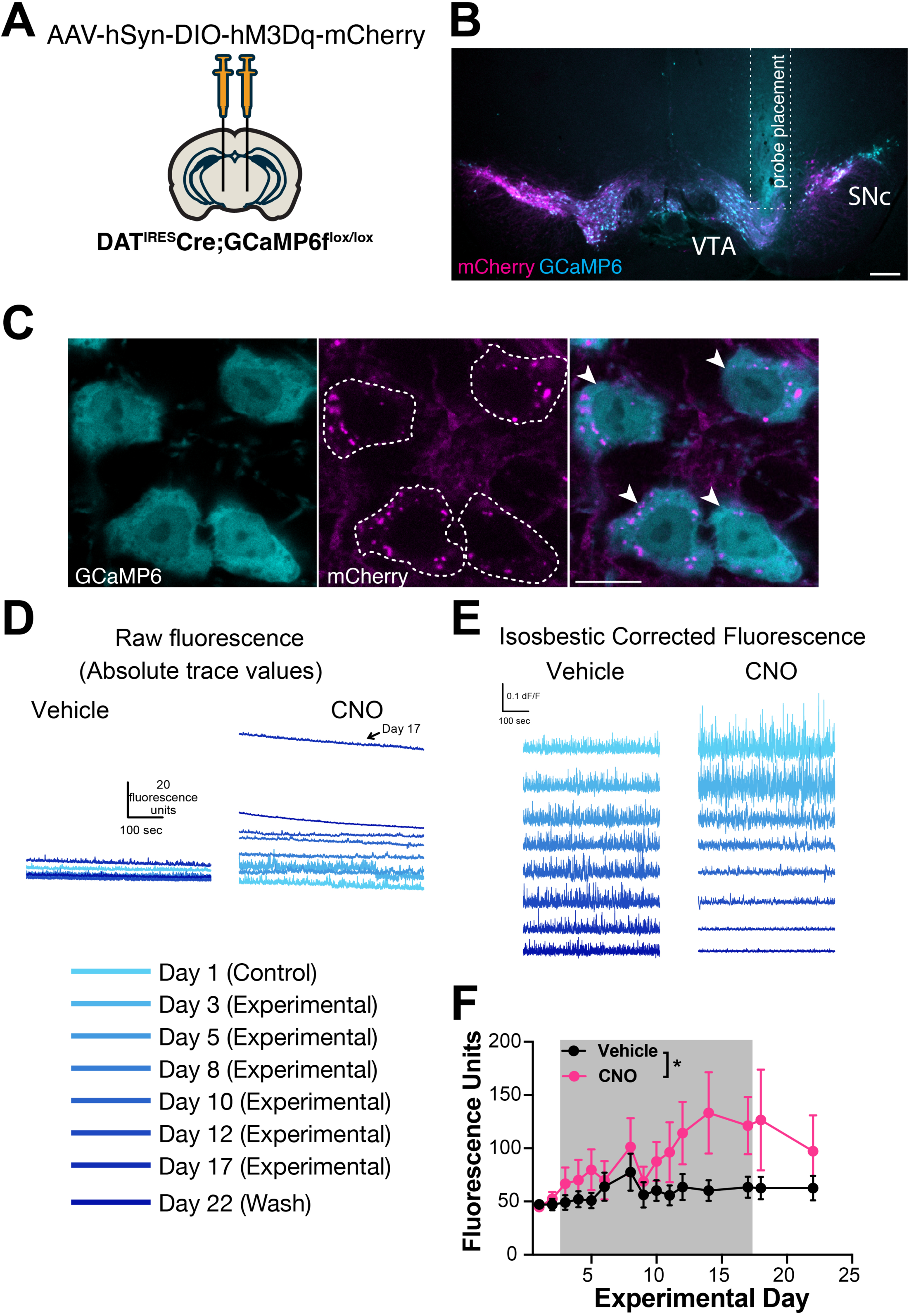
Chronic hM3Dq activation increases baseline calcium in parallel with axonal degeneration. **(A)** Transgenic mice expressing calcium-reporter GCaMP6 specifically in DA neurons were injected bilaterally with AAV-DIO-hM3Dq-mCherry and implanted with an optical probe for baseline calcium measurements during a 14-day chronic chemogenetic activation. **(B)** Representative image of photometry probe placement in mouse midbrain to record from dopamine neurons co-expressing hM3Dq-mCherry (magenta) and GCaMP6 (cyan). Scale bar is 200 μm. **(C)** Representative high-magnification images of reporter mCherry (magenta) and GCaMP6 (cyan) co-expression. Scale bar is 10 μm. **(D)** Representative raw traces of baseline calcium fluorescence in mice treated with vehicle vs CNO. **(E)** Representative isosbestic corrected traces in mice treated with vehicle vs CNO. **(F)** Baseline calcium fluorescence levels of DA neurons in mice treated with vehicle or CNO for 14 days (gray shaded area), and following wash. Error bars indicate mean ± SEM. *, p<0.05 by two-way ANOVA, n = 7 mice/group from 2 independent experiments.

Although GCaMP fluorescence is not quantitative, the consistent difference in baseline in animals tested concurrently on the same system provides confidence in our interpretation. The frequency and amplitude of calcium transients decreased with chronic CNO treatment, and during washout the frequency remained decreased while the amplitude returned to vehicle levels (Figure S4A). Notably, the decrease in frequency or amplitude is unlikely to represent a ceiling effect (i.e. where a high baseline might occlude transient detection), because while the absolute baseline and transient amplitude returned to pre-treatment levels during the washout, the frequency of transients did not. Instead, the decreases in transient frequency and amplitude may reflect less bursting activity in the neurons. As sharp peaks in genetically encoded calcium indicator signals tend to reflect bursting activity, the sustained decrease in frequency during washout may reflect more permanent dysfunction in the burst firing patterns of these neurons.

Together, these data support massive increases in DA neuron intracellular calcium over time, which may be due to hM3Dq receptor activation, calcium dysregulation, and the onset of degeneration. We also assessed time spent moving and distance traveled during photometry sessions in a subset of animals (Figure S4B). Paralleling the small initial increase in baseline calcium, there was a strong trend for CNO to increase activity over time versus controls in both measures of gross locomotion during the first week of treatment (distance traveled p=0.07, percent time spent moving p=0.06). Interestingly, the increased open field movement in the CNO group decreased back to baseline by day 11 (Figure S4B), just as calcium levels begin to climb markedly (Figure 3F), and in parallel to axonal degeneration (Figure 2A, Figure S3B).

### Identifying activation-associated transcriptomic changes with region specificity using spatial transcriptomics

To gain additional insight into the mechanisms of degeneration in this model, we used spatial genomics. We performed Visium spatial transcriptomics on midbrain and striatal sections from DAT^IRES^Cre mice that were bilaterally injected with AAV-DIO-hM3Dq-mCherry at 3 months and then received either CNO (GqCNO) or vehicle (GqVeh) drinking water for one week prior to harvest. Additional non-injected control mice also received CNO (CNO alone) (Figure 4A). Mice were harvested following one week of CNO treatment in order to quantify transcriptomic changes at a time point when some axons still remain and prior to somatic degeneration (Figure 2A, Figure S2A, Figure S5A), thereby focusing on early gene expression changes prior to neuronal death.

**Figure 4.**
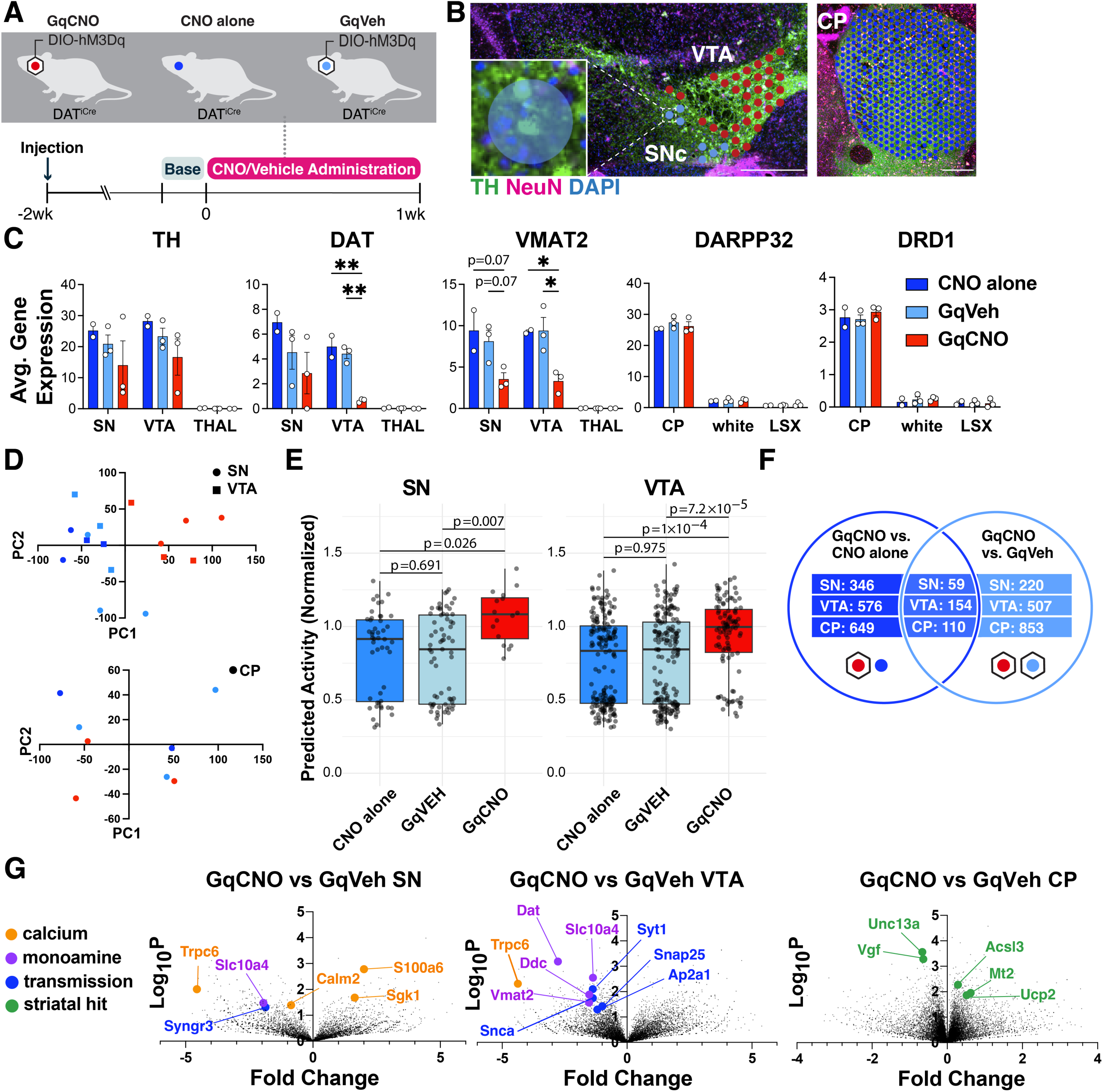
Spatial transcriptomics reveals midbrain DA and striatal target DEGs altered by chronic DA neuron hyperactivity. **(A)** DAT^IRES^Cre animals that received CNO (CNO alone, n=2 mice) or were injected with AAV-hM3Dq-mCherry and received vehicle (GqVeh, n=3 mice) or CNO (GqCNO, n=3 mice) were treated for one week before brains were flash frozen for spatial transcriptomic analysis. **(B)** Image of midbrain and striatal sections stained with TH (green), NeuN (purple), and DAPI (blue) shows discs assigned to regions of interest. Inset shows a disc containing two TH+ cell bodies. Scale bars indicate 500µm. **(C)** Expression of dopaminergic and striatal genes is confined to expected spatial regions. Expression of genes involved in DA metabolism decrease with chronic CNO. 2-49 discs were compiled per VTA and 1-7 discs were compiled per SN. 357-560 capture areas were compiled per CP. The thalamus was selected as a midbrain control region, while white matter tracts (white) and the lateral septal complex (LSX) were used as striatal controls. *p<0.05, **p<0.01, ***p<0.001 by one-way ANOVA followed by Holm-Sidak *post hoc* test in the SN, VTA, and CP. **(D)** Principal components analysis of midbrain regions (top) and the caudate putamen (bottom) for GqCNO, GqVeh, and CNO alone groups. **(E)** The deep learning NEUROeSTIMator model was used to predict neural activity of GqCNO, GqVEH, and CNO alone within the SN and VTA from Visium spatial transcriptomics. Groups were compared using KS-test (see Methods). **(F)** Hits were used for Enrichr pathway analysis if significant in both GqCNO vs CNO alone and GqCNO vs GqVeh comparisons. Gene rankings for hit analysis were established using fold change score (FCS) and signal-to-noise score (SNS). **(G)** Volcano plots comparing GqCNO vs GqVeh in the SN, VTA, and CP. Genes highlighted are also significantly altered when comparing GqCNO vs CNO alone.

We next identified 55 μm diameter barcoded discs (Visum Spatial Gene Expression) overlaying regions enriched for DA neurons within the SN and VTA in the midbrain, and the caudate putamen (CP) (Figure 4B). In the midbrain, selected discs contained at least one entire TH-positive (and mCherry-positive for injected animals) soma, and also expressed at least two of three characteristic DA genes (DAT, vesicular monoamine transporter 2 (VMAT2), or TH) above a pre- set threshold level (Figure 4C). Interestingly, DAT and VMAT2 were significantly decreased in the SN and VTA of GqCNO mice, perhaps indicating a loss of dopaminergic phenotype. In agreement with this, we found that striatal dopamine levels were markedly decreased by HPLC (Figure S6). Indeed, principal components analyses of the SN and VTA in the midbrain demonstrate that GqCNO midbrain dopaminergic regions are transcriptionally similar to one another and distinct from control groups (Figure 4D). This suggests that chronic DREADD activation induces a robust transcriptomic response in DA neurons that is independent of viral transduction or CNO administration. Consistent with this, we employed the NEUROeSTIMator method (*47*) to predict neural activity scores based on the transcriptomic assessment of the SN and VTA, finding that GqCNO samples had higher predicted activity scores than controls (GqVEH and CNO alone) (Figure 4E).

We hypothesized that chronic activation of DA neurons likely also leads to specific gene expression changes in striatal target neurons. In the striatum, discs were selected based on expression of *DARPP32* (Figure 4C). Notably, principal components analysis between GqCNO and control groups revealed less separation in the striatum than the midbrain (Figure 4D), perhaps because of the multiple inputs on striatal neurons in addition to dopaminergic axons, although future investigation is required.

We next compared the transcriptomic profiles of discs within each brain region between GqCNO and GqVeh mice, as well as in GqCNO vs CNO alone mice. Only genes that had significant differential expression between GqCNO and both control groups were assessed for pathway changes (Figure 4F). Consistent with chronic DREADD activation, pathway analyses of GqCNO vs control groups in the SN and VTA showed significant enrichment of GO Biological processes including “chemical synaptic transmission”, “synaptic vesicle exocytosis”, “positive regulation of transporter activity”, and “vesicle transport along microtubule” (Table S1). Also consistent with DREADD activation, GO Molecular Functions in the VTA included “GTP binding” and “syntaxin binding” (Table S1). Interestingly, both CP and SN GO Molecular Functions included “calcium channel regulator activity”, suggesting a larger circuit-level change in calcium channel regulation as a result of chronic DREADD activation (Table S1).

Consistent with this, there were a number of differentially expressed genes in the SN and/or VTA that regulate neural transmission, calcium, and activity (Figure 4G, Figure S5B, S5C, Figure S7, Figure S8). These included increased expression of S100a6, a calcium sensor and modulator (*48*) and Sgk1, a serine/threonine kinase regulated by intracellular calcium involved in the regulation of ion channels, membrane transporters, cellular enzymes, and transcription factors (*49*). Interestingly, Sgk1 expression trended up in the CP, supporting circuit-level changes in calcium signaling. Increased S100a6 expression has been observed in AD and ALS (*50*), while Sgk1 upregulation has been associated with DA neuron death in rodent toxin models of PD (*51, 52*). Conversely, the expression of calmodulin-2 (Calm2), which mediates control over a large number of enzymes and ion channels as a result of calcium binding (*53*), and Trpc6, which is thought to form a non-selective calcium-permeant ion channel (*54*) were both decreased. Decreased Trpc6 may be a homeostatic response to chronically high intracellular calcium. These changes in calcium-regulating genes are consistent with the changes in baseline intracellular calcium observed in DA neurons by GCaMP photometry (Figure 3), and may reflect early processes that either cause or contribute to the onset of degeneration, or alternatively compensate to protect against degeneration.

Pathway-level changes in transmission were driven by decreases in several synaptic-vesicle genes, including: synaptotagmin 1, a calcium sensor that triggers transmitter release including striatal dopamine release (*55*); Snap25, a t-SNARE that regulates transmitter release (*56*); Ap2a1, a component of the adapter protein complex AP-2 (*57*); and Syngr3, a synaptic vesicle-associated protein that may regulate DAT function (*58*). Interestingly, α-synuclein was also significantly downregulated in the VTA. α-synuclein is enriched in presynaptic terminals, and has been shown to modulate DA release (*59*). Together, these results suggest that chronic DREADD-activation of DA neurons results in significant changes in synaptic transmission which may result in altered calcium regulation and DA release.

To begin to interrogate the role of calcium in hyperactivity-induced degeneration, we performed a pilot experiment in which mice injected with saline in the left midbrain and AAV- hM3Dq(DREADD)-mCherry in the right midbrain were then implanted subcutaneously with slow-releasing placebo or isradipine pellets (Figure S9A). Mice were not acutely tested for responsiveness to CNO, as wheel running data is less clear with unilateral manipulation. Instead, DREADD expression was validated *post hoc* by immunofluorescence, and animals with little or no DREADD expression were excluded from further analysis. Isradipine blocks Ca_v_1.2 and Ca_v_1.3 channels that are highly expressed in SNc neurons (*60, 61*). Isradipine treatment has been shown to return SNc DA neurons to a juvenile form of pacemaking, and chronic treatment has been shown to reduce calcium-dependent mitochondrial oxidant stress (*60, 62*). To confirm the presence of systemic isradipine, we measured isradipine levels in blood plasma via HPLC (Figure S9B). Following two weeks of vehicle or CNO administration with placebo or isradipine release, we assessed striatal TH immunoreactivity and mCherry immunofluorescence. Consistent with previous results, we observed decreases in TH and mCherry in CNO-treated AAV-Gq hemispheres, but chronic isradipine administration did not rescue this loss (Figure S9C, S9D). Notably, the absence of rescue may result from a lack of statistical power due to small sample size, or due to the presence of non-responders. Interestingly, chronic isradipine induced some decrease in TH immunoreactivity in vehicle-treated AAV-Gq hemispheres without reductions in mCherry, suggesting that chronic isradipine may induce TH downregulation without degeneration.

We also identified differentially expressed genes (DEGs) in the striatum. Upregulated genes included: Mt2, a metallothionein protein that acts as an anti-oxidant to protect against free radicals (*63*); Acsl3, an acyl-CoA synthetase that stimulates fatty acid synthesis (*64*) and could supply energy substrates; and the mitochondrial uncoupling protein Ucp2. Downregulated genes included: Unc13a, which is involved in neurotransmitter release via vesicle maturation and has been implicated in ALS (*65, 66*), and Vgf, a secreted growth factor whose expression is also decreased in PD (*67*).

We also evaluated the SN DEGs (n=59) and additional genes of interest identified within our DREADD mouse model with gene expression changes within human control and early PD subjects (Figure S10A,B). We employed the Nanostring GeoMx assay to assess whole transcriptome changes of dopamine neurons (TH+ masked, see Methods) within the SNc ventral tier of Controls and Early PD subjects. Of the 59 DEGs, 51 were found within the human dataset, and these genes exhibited broadly congruent directional changes between mouse and human (Chi-square p-value = 0.05). This suggests that similar mechanisms may be at play in the PD midbrain and the DREADD mouse model. To investigate this further, we considered genes that emerged from the mouse dataset involved in calcium, transmission, and dopamine, and evaluated them in the early PD samples (Figure S10B, S10C). We found that expression of Syt1, Syngr3, and Calm2 were all decreased in both mouse and human datasets. Similarly, we found that expression of Slc18a2 (VMAT2) and Slc10a4 were also both decreased in the mouse and human data. These findings may suggest common mechanisms of calcium dysregulation and altered dopamine metabolism in an early stage of degeneration in our mouse model and in human PD. Additionally, differential gene expression analysis revealed that HSPA4 and EIF3G were downregulated (FDR < 0.02) within the SNc ventral tier DA neurons of early PD subjects compared to controls and in the mouse model (Figure S10A) highlighting other potential common pathogenic mechanisms. Additional investigation is required to determine the functional impact of these genetic changes on dopamine neuron and dopamine neuron subtype function and vulnerability.

To further assess these findings on a single cell basis, we performed high resolution Xenium spatial transcriptomics on adjacent sections from the same tissues (Figures S11). Sections processed for Xenium were then processed for mCherry immunofluorescence. Expression of mCherry, along with anatomy, and expression of characteristic dopaminergic genes (TH, DAT, VMAT2) were used to select AAV-transduced dopaminergic nuclei in the SNc and VTA. In these nuclei, TH and VMAT2 expression levels were similar across groups, but expression of DAT was decreased in GqCNO nuclei. Similar to the lower-resolution Visium, expression of Slc10a4 was also decreased in GqCNO nuclei. Interestingly, other markers of vulnerable SNc neurons were also decreased in GqCNO SNc nuclei, including Aldh1a1, Sox6, and Anxa1 (*68–70*). Downregulation of Aldh1a1 has also been reported in PD (*71*). These subtype markers cannot be assessed in the lower-resolution Visium dataset due to their relative scarcity and the large size of the Visium capture areas, but their differences are appreciable with the single-nucleus resolution of Xenium. Together, these results support a loss of a dopaminergic phenotype, especially in vulnerable nigral dopaminergic nuclei.

## Discussion

Here we establish a new model system to chronically activate midbrain DA neurons, and show that this prolonged activation leads to the preferential and step-wise degeneration of SNc DA neurons, starting with loss of terminals and progressing to neuronal death.

### New model to chronically activate DA neurons

Prior studies have activated midbrain DA neurons by both optogenetics and DREADDS to study the impact of DA neuron activity on sleep (*72*) and the spread of pathological α-synuclein (*73*). However, in order to accurately model the impact of neural activity on neurodegeneration, we hypothesized that activity must be chronically increased. Unlike other studies which have used once or twice daily injections of CNO, we delivered CNO in the drinking water to ensure a more continuous, sustained impact on neural activity. We confirmed that chronic 1, 2, or 4 week hM3Dq activation results in sustained increases in intracellular calcium levels and functional behavioral readouts, suggesting that our model induces long-lasting changes in dopamine neuron activity. Moreover*, ex vivo* electrophysiology demonstrated that the basal level of DA neuron activity does indeed increase with CNO treatment. An important question, however, is whether our model accurately recapitulates how neural activity increases in physiologic or pathologic situations.

Compensatory increased or decreased rates of DA neuron pacemaking may help to maintain striatal DA as it is depleted in PD, or to protect against energy failure by decreasing energy consumption. Meanwhile, changes in bursting may reflect circuit-level adaptations to PD in response to ongoing degeneration. Irregular or silenced pacemaking is commonly observed in toxin-based models of PD (*74–78*), while increased induced bursting is observed after massive loss of DA neurons or complete loss of complex I (*1, 6–8*). These changes could further promote the degeneration of remaining DA neurons. As such, in future studies it will be important to understand how changes in the pattern of activity (i.e. bursting versus pacing) impact degeneration. In other PD models, DA neuron activity is altered prior to degeneration (*9, 12, 13*). The energetic demands of neural activity position altered activity as a possible contributor to the degenerative process in energetically vulnerable DA neurons, either as an initiating force or as a driver of disease progression.

One possible outcome of chronic hyperactivation is depolarization block (DB), a process that occurs even in healthy neurons wherein prolonged excitation induces persistent depolarization and blockade of spike generation (*79*). DB occurs in midbrain DA neurons following chronic treatment with antipsychotic drugs (*80*), and DB in SNc neurons specifically is associated with extrapyramidal symptoms (*81, 82*). Lesions of DA neurons with 6-OHDA can also increase susceptibility to DB (*83*). In our chronic hyperactivation model there is a strong decrease in dark cycle wheel running behavior that may be consistent with DB (Figure 1C, Figure S1B). Prolonged excitation along with progressive degeneration in our model may put DA neurons at greater risk of entering DB. However, CNO-treated animals have increased rather than decreased open field movement (Figure S4B). Our NEUROeSTIMator data also predicts higher activity scores in GqCNO samples compared to GqVEH and CNO alone controls (Figure 4E). Moreover, while DB can’t be assessed in slices due to its reliance on afferent input, we observed increased pacemaking in CNO-treated neurons, further arguing against prominent induction of DB. Given that VTA and SNc neurons can have differential susceptibilities to DB in response to antipsychotics, it is possible that a subpopulation of DA neurons in our model periodically enter DB without gross effects on locomotion. Additional data such as *in vivo* electrophysiology will be necessary to fully investigate the possibility of DB.

As expected, chronically increasing neural activity increased the movement of mice during the light cycle. However, it was notable that movement as measured by wheel use in the home cage did not increase as robustly as did movement in the open field chamber (Figure S4B), perhaps indicating that the impact on movement is greater in novel environments. This may reflect the effects of midbrain DA neurons on motivation and exploration (*44, 84*). The later return to control levels of locomotion midway through CNO treatment may indicate the onset of synaptic dysfunction or degeneration. In addition, our electrophysiologic analyses focused on cell body function, but the loss of terminals starting by ≈1 week (Figure S3B) raises the possibility that synaptic function might be disrupted even earlier, in parallel to behavioral changes. Meanwhile, chronically increasing activity unexpectedly and robustly decreased movement during the dark cycle (when mice should be most active) within 3-4 days of starting CNO (Figure 1C, Figure S1B). One possibility is decreased dark cycle movement results from a disruption in sleep, which might occur given the mild increased use of running wheels during the light cycle. Indeed, prior studies show that acute inhibition or stimulation of VTA DA neurons by optogenetics or DREADDs suppresses or promotes wakefulness, respectively (*72, 85*). These data support a critical role for VTA DA neurons in maintaining wakefulness, and suggest that increased activity might promote wakefulness in our activity model. Future investigations assessing the impact of chronic SNc versus VTA DA neuron activation on circadian rhythms and sleep are needed to determine if these DA neurons similarly influence sleep in the setting of prolonged stimulation, and how potential sleep changes might contribute to the effects of chronically increased neural activity on motor function.

### Chronic chemogenetic activation drives degeneration

How does increasing the activity of SNc and VTA DA neurons impact these neurons, as well as their striatal targets? Analysis by spatial transcriptomics near the onset of axonal degeneration reveals a clear decrease in the expression of genes involved in DA synthesis (TH), uptake (DAT) and storage (VMAT2) in the SNc and VTA, perhaps reflecting neuronal stress that precedes degeneration or an attempt to decrease DA release. Further investigation will be needed to determine whether these gene expression changes are associated with functional deficits. Decreases in presynaptic transmission-related genes (Syt1, Snap25, Snca, Ap2a1) in midbrain dopamine neurons are consistent with axonal dysfunction and loss that are observed later in CNO treatment. Moreover, consistent with the role of hM3Dq DREADDs in increasing cytosolic calcium (*46*) on a systems level, we observed a significant change in expression of genes involved in the regulation of calcium channels (Supplemental Table 1), as well as serum and glucocorticoid-inducible kinase 1 (Sgk1), a protein-serine/threonine kinase activated by calcium (*86*), in both the midbrain and striatum. Our photometry analysis also raise the possibility of a central role for calcium in activity-induced degeneration, as calcium levels in midbrain DA levels rose markedly following 7 days of CNO (Figure 3F), as axonal degeneration progresses (Figure S3B).

The mechanism of preferential degeneration remains to be determined, but is likely triggered by increased cytosolic calcium levels associated with hM3Dq activation (*46*). SNc DA neurons are believed to be intrinsically vulnerable to increased calcium due to their pacemaking activity driven by CA_V_1.3 channels (*60*), combined with the high energetic requirements of removing calcium from neurons and lack of calbindin expression (*33*). Further investigation is required to determine if DREADDs differentially increase calcium levels in SNc vs VTA DA neurons, and/or if SNc DA neurons are more vulnerable to the same level of elevation. Further, it remains unknown whether increased calcium leads directly to degeneration, for instance by triggering energy failure or increasing ROS, or if it indirectly causes neurodegeneration by triggering potentially toxic downstream processes such as increased cytosolic or extracellular DA release.

Excessive glutamate leads to large influxes of calcium that can trigger excitotoxicity leading to neurodegeneration and cell death (*87*). Chronic, mild calcium elevation can also dysregulate mitochondrial calcium buffering, and increase sensitivity to cytosolic calcium toxicity (*39*), and the route of calcium entry can also impact toxicity (*88*). Isradipine, a blocker of Ca_v_1.2 and Ca_v_1.3 channels, failed to protect against hyperactivity-induced degeneration in our model (Figure S9). However, it is possible that other interventions against calcium-induced toxicity such as inhibition of NMDA receptors or the mitochondrial calcium uniporter may be protective. Alternatively, degeneration could be driven by the toxicity of excessive extracellular dopamine as a consequence of elevated activity. Future studies will explore these potential mechanisms.

If increased DA neuron activity contributes to degeneration in PD, then one might also expect gene expression changes in our model to mirror those seen in PD. Indeed, we found that mouse DEGs exhibited broadly congruent directional changes in the human dataset (Figure S10). This raises the possibility that similar mechanisms of degeneration and adaptation may be at play in both our mouse model and in PD. Interestingly, HSPA4 was significantly downregulated in the mouse and human datasets. Heat shock proteins broadly function to remove misfolded proteins, but the function of HSPA4 in neurons is not well described. However, in rat neural stem cells, HSPA4 is upregulated by selegiline, a type B monoamine oxidase inhibitor used to treat PD, and HSPA4 upregulation reduces ROS levels and mitochondrial DNA damage following hydrogen peroxide exposure (*89, 90*). In MPTP-treated mice, deacetylation of HSPA4 is linked to decreased microglial activation and neuroinflammation (*91*). In our activity model, decreased HSPA4 expression might indicate a failure to respond to inflammation, promoting neurodegeneration. Further studies will be required to investigate the role of HSPA4 and inflammation in chronic hyperactivation-induced toxicity.

Notably, we compared our mouse transcriptomic data at a stage when there is mild terminal loss and no somatic degeneration with early human PD patient tissues where there is limited dopamine neuron loss, raising confidence in the relevance of the comparison. However, it is important to consider that our Visium transcriptomic analysis has limited cell type resolution and our high resolution Xenium analysis is limited to nuclear transcripts. Therefore, important areas of future investigation include higher spatial resolution in the analysis of mouse tissue, and assessment of additional disease stages, brain areas, and cell types in human PD patient tissues.

In some cases, disease proteins or circuit changes may decrease neural activity. As such, it will be important to understand in which contexts activity increases or decreases, and also how decreasing DA activity influences degeneration. Indeed, if activity does promote degeneration, it will also be important to understand if decreasing DA neuron activity is protective, and if activity can be decreased or modulated without compromising motor function. Interestingly, chronic nicotine, which may mediate the protective association of smoking in PD (*92, 93*), inhibits SNc DA neurons through agonism of nicotinic acetylcholine receptors expressed on presynaptic GABAergic terminals (*94*). In addition, a proposed mechanism of action for the beneficial effects of deep brain stimulation (DBS) in PD is the inhibition of the subthalamic nucleus. As such, it will also be important to know how DBS influences DA neuron activity and if it has the potential to influence neurodegeneration. Interestingly, recent clinical trial data raise the possibility that DBS might slow disease progression when administered to early-stage patients (*95*).

In summary, our data show that chronically increasing the activity of DA neurons can produce toxicity, and that SNc neurons may be more susceptible to this effect than VTA neurons. Considering that DA neuron activity may increase to compensate for other dying DA neurons, or in response to certain disease proteins, our data support the hypothesis that increased neural activity contributes to the pathophysiology of at least a subset of PD. Proving this, and determining how this evolves with disease stage, will be important goals for future research.

## FIGURE LEGENDS

**Figure S1. Additional behavior data.**

**(A)** (Left) Representative running wheel traces of a DAT^IRES^Cre mouse injected with AAV-hM3Dq(DREADD)-mCherry following i.p. injection of either saline (left) or CNO (right). Dotted line indicates time of injection. (Right) Average running wheel rotations per minute over 3-hour time period of DAT^IRES^Cre mice injected with AAV-hM3Dq(DREADD)-mCherry following saline or CNO i.p. injection.

**(B)** Average running wheel rotations per minute over 12-hour light (top) and dark (bottom) cycles for two independent cohorts of DAT^IRES^Cre mice injected with AAV-hM3Dq(DREADD)-mCherry. CNO (300 mg/L) or vehicle (2% sucrose in water) was administered *ad libitum* via drinking water for two weeks and the animals perfused the next day. Some data was lost from days 2-6 and 13-14 due to technical issues during Cohort 2. Open circles in light cycle denote incomplete datasets (3hrs). n=5 animals/group.

**(C)** (Left) Average running wheel rotations per minute over 12-hour light (top) and dark (bottom) cycles for non-injected DAT^IRES^Cre (CNO alone) mice. (Right) Mean wheel usage for selected days during the experiment, segregated by light (top) or dark (bottom) cycles. n=5 animals/group.

**(D)** (Left) Average running wheel rotations per minute over 12-hour light (top) and dark (bottom) cycles for DAT^IRES^Cre mice injected with AAV-hM3Dq(DREADD)-mCherry that did not display acute running wheel responses to i.p. of CNO. (Right) Mean wheel usage for selected days during the experiment, segregated by light (top) or dark (bottom) cycles. n=5 animals/group. *p<0.05 by two-way ANOVA followed by Holm-Sidak *post hoc* test.

**Figure S2. Additional electrophysiology data.**

**(A)** SNc DA neurons expressing the DREADD and activated for one week *in vivo* showed alterations in CNO responses and overall physiology. *I*_h_ magnitude was measured in whole cell voltage clamp configuration, V_holding_ = −60 stepping to −120 mV in neurons from vehicle-treated cells vs CNO-treated cells. Firing regularity and input resistance were measured during the first 2 min of whole cell recordings. Spontaneous AP waveforms were compared across groups for threshold membrane potential, peak membrane potential, and duration. Initial membrane potential was measured upon first breaking into the cells for whole cell recordings. ** p ≤ 0.01, *** p ≤ 0.005 by t-test or permutation (non-parametric) analysis.

**(B)** VTA DA neurons expressing the DREADD and activated for one week *in vivo* showed alterations in CNO responses and overall physiology. Time course of responses to bath application of 1 μM CNO *ex vivo* measured in current clamp in neurons from vehicle-treated and CNO-treated mice. Spontaneous firing rate was measured during the first 2 min of whole cell recordings. *I*_h_ magnitude was measured in whole cell voltage clamp configuration, V_holding_ = −60 stepping to −120 mV in neurons from vehicle-treated cells vs CNO-treated cells. Firing regularity and input resistance were measured during the first 2 min of whole cell recordings. Spontaneous AP waveforms were compared across groups for threshold membrane potential, peak membrane potential, and duration. Initial membrane potential was measured upon first breaking into the cells for whole cell recordings. * p ≤ 0.05 by t-test or permutation (non-parametric) analysis.

**(C)** Example traces of spontaneous firing activity from whole cell current clamp recordings (I = 0 pA) in SNc and VTA neurons from vehicle and CNO treated mice.

**(D)** The traces from each neuron tested with 1 μM CNO. These data are summarized in Figure 1E.

**Figure S3. Additional histological data.**

**(A)** (Top) Example images of TH immunoreactivity in striatal sections of non-responder mice treated for two weeks with vehicle (left) or CNO (right) via drinking water. DA neuron projection areas in dorsal and ventral striatum are indicated with dotted lines. Quantification for TH optical density shows preferential loss in CPu. n = 5 animals/group, 2-4 sections/animal. Scale bar indicates 200μm. Error bars indicate mean ± SEM. Analyzed by two-way ANOVA and Holm-Sidak *post hoc* test.

**(B)** Example images of TH immunoreactivity in striatal sections of hM3Dq-expressing mice treated for one week with vehicle **(left)** or CNO **(right)** via drinking water. DA neuron projection areas in dorsal and ventral striatum are indicated with dotted lines. Quantification for TH optical density shows a trend for preferential loss in CPu. n = 5 animals/group, 3-5 sections/animal. Scale bar indicates 200μm. Error bars indicate mean ± SEM. Analyzed by two-way ANOVA and Holm-Sidak *post hoc* test. CPu: caudate putamen, NAc-C: nucleus accumbens core, NAc-Sh: nucleus accumbens shell, OT: olfactory tubercule.

**Figure S4. Additional photometry data.**

**(A)** Calcium transient frequency (left) and amplitude (right) in DA neurons in hM3Dq-expressing GCaMP6f^fl^ mice treated with vehicle or CNO for 14 days (gray shaded area), and following wash. Error bars indicate mean ± SEM. **p<0.01 by two-way ANOVA, n = 7 mice/group from 2 independent experiments.

**(B)** Distance traveled (left) and percent time spent moving (right) during photometry sessions in mice treated with vehicle or CNO for 14 days (gray shaded area) and following wash. Error bars indicate mean ± SEM. N = 3-4 mice/group. Analyzed by two-way ANOVA.

**Figure S5. Additional spatial transcriptomic data following chronic hyperactivation.**

**(A)** Anti-TH immunofluorescence of striatal sections processed for spatial transcriptomics demonstrate loss of TH+ fibers in GqCNO sections after one week of chronic hyperactivation (white arrows). Scale bars indicate 1mm.

**(B)** Volcano plots comparing GqCNO vs CNO alone in the SN, VTA, and CP.

**(C)** Bar graphs of individual midbrain and striatal genes of interest. N=2-3 animals/group. Error bars indicate mean ± SEM. *p<0.05, **p<0.01, ***p<0.001 by one-way ANOVA followed by Holm-Sidak *post hoc* test.

**Figure S6. Striatal DA is decreased with chronic activation.**

HPLC of CPu dissected from fresh-frozen brain tissue from DAT^IRES^Cre mice bilaterally injected with AAV-hM3Dq(DREADD)-mCherry and treated with vehicle or CNO water for one week (GqVEH and GqCNO) or uninjected DAT^IRES^Cre treated with CNO water for one week (CNO alone). Error bars indicate mean ± SEM. N = 2-3 mice per group. ***p<0.001 by two-way ANOVA followed by Holm-Sidak *post hoc* test. NE: norepinephrine, DOPAC: 3,4-dihydroxyphenylacetic acid, DA: dopamine, 5HIAA: 5-hydroxyindoleacetic acid, HVA: homovanillic acid, 5HT: 5-hydroxytryptamine or serotonin, 3MT: 3-methoxytyramine.

**Figure S7. Top differentially expressed genes in the mouse SN and VTA.**

Heatmaps of the top differentially expressed SN and VTA genes, expressed as the log_2_ fold score.

**Figure S8. Top differentially expressed genes in the mouse CP.**

Heatmaps of the top differentially expressed CP genes, expressed as the log_2_ fold score.

**Figure S9. Chronic isradipine treatment does not rescue axonal degeneration.**

**(A)** Experimental paradigm. DAT^IRES^Cre mice were injected in the left midbrain with saline and the right midbrain with AAV-hM3Dq(DREADD)-mCherry and allowed to recover for 2 weeks. Mice were then subcutaneously implanted with either placebo or isradipine slow-release pellets (3μg/g/d) and started either vehicle or CNO administration for 2 weeks.

**(B)** Isradipine plasma concentration at the end of the experiment. N = 9 animals/group.

**(C)** (Top left) Example images of TH immunoreactivity in striatal sections of mice implanted with placebo or isradipine pellets and treated for two weeks with vehicle or CNO via drinking water. (Top right) Quantification for TH optical density in the CPu shows loss with CNO treatment in hM3Dq-injected hemispheres that is not rescued with chronic isradipine treatment. N = 5-9 hemispheres per group, 2-3 sections/animal. Scale bar indicates 200μm. Error bars indicate mean ± SEM. ***p<0.001 by three-way ANOVA followed by Holm-Sidak *post hoc* test. p=0.006 for the interaction between CNO treatment and hM3Dq injection by three-way ANOVA. (Bottom left) Example images of mCherry immunofluorescence in striatal sections of mice implanted with placebo or isradipine pellets and treated for two weeks with vehicle or CNO via drinking water. (Bottom right) Quantification for mCherry fluorescence density in the CPu shows loss with CNO treatment in hM3Dq-injected hemispheres that is not rescued with chronic isradipine treatment. N = 5-9 hemispheres per group, 2-3 sections/animal. Scale bar indicates 200μm. Error bars indicate mean ± SEM. ***p<0.001 by three-way ANOVA followed by Holm-Sidak *post hoc* test. CPu: caudate putamen.

**Figure S10. Assessment of mouse DEGs in human PD and control SNc samples.**

Heatmap of normalized gene expression (per ROI) values from GeoMx spatial transcriptomic analysis of human SNpc ventral tier dopamine neurons (TH+) within **(A)** 51 DEGs and **(B)** additional genes of interest identified within the mouse model. N = 10 control and 8 early PD samples. ‘Limma voom’ methodology was used to assess differential gene expression between control and PD samples. PMD, post-mortem delay; DV200, RNA integrity number equivalent. **(C)** Bar graphs of individual midbrain genes of interest in mouse and PD samples.

**Figure S11. Xenium spatial transcriptomics.**

Images show anti-mCherry immunofluorescence in a midbrain section processed for Xenium spatial transcriptomics. Sections used for Xenium are adjacent to those used for Visium. Inset includes transcripts for TH, DAT, and VMAT2. Scale bars indicate 500μm (left) and 50μm (right). Quantifications for average nuclear counts for individual transcripts of interest. N = 2-3 mice per group. **p<0.01 by two-way ANOVA followed by Holm-Sidak *post-hoc* test.

**Supplementary Table S1. Pathway analyses for the VTA, SN, and CP following chronic hyperactivation.**

Gene Ontology Molecular Function and Biological Process terms for differentially expressed genes in the VTA, SN, and CP generated with the Enrichr webtool.

**Supplementary Table S2. Demography of the human post-mortem cohort assayed by GeoMx.**

Values are presented as median (IQR). The comparison of Age at death, Post-mortem delay and DV200 between groups was made using the Welch Two Sample t-test. The comparison of gender between groups was made using chi-square test. * P < 0.05, ** P < 0.001.

## Methods

### Experimental model and subject details

#### Mice

Mice were group-housed in a colony maintained with a standard 12-hour light/dark cycle and given food and water ad libitum. All mice received food on the cage floor. All animal experimental procedures were conducted in accordance with the Guide for the Care and Use of Laboratory Animals, as adopted by the National Institutes of Health, and with approval from the University of California, San Francisco Institutional Animal Care and Use Committee. All mice were housed in a state-of-the-art barrier facility managed by the UCSF Laboratory Animal Resource Center (LARC). Animal care and use in this research are covered under the UCSF “Assurance of Compliance with PHS Policy on Humane Care and Use of Laboratory Animals by Awardee Institutions” number A3400-01. Experiments were performed on age-matched mice. All mouse lines were maintained on a C57Bl/6 background (The Jackson Laboratory; RRID:IMSR_JAX:000664). DAT^IRES^Cre mice (*96*) were obtained from the Jackson laboratory.

### Methods

#### Chemogenetics

A detailed protocol for mouse stereotaxic surgery can be found at https://dx.doi.org/10.17504/protocols.io.b9kxr4xn. DAT^IRES^Cre homozygous mice (RRID:IMSR_JAX:006660) were injected bilaterally with 1 μL of rAAV8-hSyn-DIO-hM3Dq-mCherry (UNC Vector Core, AddGene, RRID: Addgene_50459) into the midbrain (−3.0 mm anterior-posterior, ±1.2 mm medial-lateral, −4.3 mm dorsal-ventral) using a stereotaxic frame (Kopf) and a microliter syringe (Hamilton).

A detailed protocol for monitoring mouse activity can be found at https://dx.doi.org/10.17504/protocols.io.3byl4qqe8vo5/v1. Two weeks following surgery, animals were single-housed and habituated to wireless running wheels (MedAssociates ENV-047, RRID: SCR_024879) connected to a hub (MedAssociates DIG-807, RRID: SCR_024880,) for locomotion recordings and water bottles (Amazon) for drinking. A detailed protocol for validating responsiveness of DREADD-injected DAT^IRES^Cre mice to CNO can be found at https://dx.doi.org/10.17504/protocols.io.6qpvr3yrpvmk/v1. Validation of responsiveness was done for all DREADD experiments, except for those unilaterally injected for chronic isradipine treatment (Figure S9). All mice had access to a running wheel during their treatment, except those for fiber photometry and chronic isradipine treatment. The mice used for photometry underwent 7 days of running wheel access approximately 3 weeks prior to the beginning of the experiment. The photometry headcaps sterically prevented mice from having access to a running wheel in their home cage. Mice used for chronic isradipine treatment were unilaterally injected with hM3Dq which does not lend to clear interpretation of running wheel data.

A detailed protocol for preparing and administering CNO via drinking water can be found at dx.doi.org/10.17504/protocols.io.n2bvj33oblk5/v1. CNO (NIMH, Tocris 4936) was administered ad libitum in 2% sucrose water at 300mg/L. CNO and vehicle waters were made fresh weekly and stored at 4°C protected from light.

#### Ex Vivo Recording

A detailed protocol for ex vivo electrophysiology can be found at dx.doi.org/10.17504/protocols.io.261gedqn7v47/v1. DAT^IRES^Cre mice injected bilaterally with rAAV8-hSyn-DIO-hM3Dq-mCherry and chronically treated with vehicle or CNO for one week were provided to the electrophysiologist blind to in vivo treatment. Mice were deeply anesthetized with isoflurane, decapitated with a rodent guillotine, and the brains were removed. Horizontal brain slices (150 um thick) containing the SNc were prepared using a Vibratome (Campden Instruments,7000smz-2). Slices were cut in ice cold aCSF solution containing (in mM): 119 NaCl, 2.5 KCl, 1.3 MgSO_4_, 1.0 NaH_2_PO_4_, 2.5 CaCl_2_, 26.2 NaHCO_3_, and 11 glucose saturated with 95% O_2_–5% CO_2_ and allowed to recover at 33°C in aCSF for at least 1 hr.

Slices were visualized under an *Axio Examiner A1* equipped with Dodt and IR optics, using a Zeiss Axiocam 506 mono and Neurolucida 2023 (MBF Biosciences) software. Neurons were identified as DREADD-expressing prior to patching with fluorescent imaging of mCherry expressed concurrently with DREADDs. Whole-cell patch-clamp recordings were made at 33°C using 3– 5 MOhm pipettes containing (in mM): 123 K-gluconate, 10 HEPES, 0.2 EGTA, 8 NaCl, 2 MgATP, and 0.3 Na_3_GTP, pH 7.2, osmolarity adjusted to 275. Biocytin (0.1%) was also included in the internal solution to identify neurons after recordings where desired.

Recordings were made using Sutter IPA and SutterPatch v2.3.1 software (Sutter Instruments), filtered at 5 kHz and collected at 10 kHz. Liquid junction potentials were not corrected. *I*_h_ was recorded by voltage clamping cells at −60 mV and stepping to −40, −50, −70, −80, −90, −100, −110, and −120 mV. Cells were recorded in current-clamp mode (*I=* 0 pA) for measuring spontaneous firing rates and CNO responsivity. For CNO testing, spontaneous firing rate or resting membrane potential were monitored until a stable baseline was observed for at least 5 min. Then perfusion solution was switched 1 uM CNO for 5 min.

When recordings were completed, slices were drop fixed in 4% formaldehyde in PBS for at least 2 hr.

For quantifications, spontaneous firing rate was measured as the mean firing rate during the first 120 sec of whole cell recording. Once every 10 sec a brief hyperpolarizing pulse was applied, and the input resistance of the neuron was quantified from this test, averaged across the measurements made during the first 2 min of recording. AP waveform measurements were made from averages across at least 8 APs from this recording interval. *I*_h_ magnitude was quantified as the difference between the initial steady state response to the −120 mV step and the asymptote of the slow current sag.

Statistical analyses were completed in R (4.2.3), first testing whether data met the criteria for parametric statistical evaluation. Those datasets that met criteria were compared by unpaired Student’s t-test. Those that did not meet these criteria were compared by permutation test. Code for the electrophysiology analysis can be found on GitHub at https://github.com/eb-margolis-neuroscience-lab/R-general-Margolislab/blob/main/permtestsEBM.R.

All salts and reagents were purchased from Sigma except CNO (Tocris).

#### Ex Vivo Data Analysis

Results are presented as mean ± S.E.M or with kernel density estimations. All but 1 neuron recorded in current clamp was quiescent during CNO testing. This neuron both depolarized and increased firing rate in response to CNO and therefore was included in the time course average figure. For all data parametric assumptions were tested to choose between t-test (parametric) or permutation (non-parametric) analysis.

### Fiber photometry

A detailed protocol for collecting fiber photometry data can be found at dx.doi.org/10.17504/protocols.io.bp2l6xwxzlqe/v1. 3-month-old Ai148 mice (RRID:IMSR_JAX:030328) bred with DAT^IRES^Cre were injected with rAAV8-hSyn-DIO-hM3Dq-mCherry as described above. After allowing 3 weeks for expression of the hM3Dq construct, locomotion was tested using IP injection of CNO to determine robust hM3Dq expression. Mice that exhibited increased locomotion following CNO injection were then implanted with optical fibers (400 µM, 0.48 NA). Mice were administered vehicle or CNO water for 14 days.

*In vivo* calcium data was acquired using a custom-built photometry system. An RZ5P fiber photometry processor (TDT, RRID: SCR_024878) and Synapse software (TDT, RRID: SCR_006307) were used to control LED output and acquire the photometry signal. Using this system, two LEDs were used to control GCaMP and isosbestic excitation (470 nm and 405 nm, respectively, Thorlabs). LEDs were sinusoidally modulated at 211 Hz (470 nm) and 531 Hz (405 nm) and entered a four-port fluorescence mini cube (Doric Lenses). The combined output was coupled to a fiber-optic patch cord (400 µm, 0.48 NA, Thorlabs), which then mated to the fiberoptic cannula in the mouse brain. The emitted light was collected onto a visible femtowatt photoreceiver module (AC low, Newport) and sampled at 60 Hz. Photometry data was then extracted via proprietary TDT software using MATLAB (MathWorks, RRID: SCR_001622). Code for the fiber photometry analysis can be found on GitHub at https://github.com/yoshitakasei/kreitzerPhotometry.

### Open Field Behavior

A detailed protocol for open field analysis can be found at dx.doi.org/10.17504/protocols.io.36wgq33rklk5/v1. Spontaneous locomotor activity was assessed while simultaneously recording fiber photometry data. Videos acquired during photometry sessions were analyzed using Ethovision software (Noldus, RRID: SCR_000441) to calculate total distance traveled and percent of time spent moving.

### Chronic Isradipine Treatment and Plasma Detection

DAT^IRES^Cre mice were unilaterally inject in the right midbrain with AAV-hM3Dq as detailed above and in the left midbrain at the same coordinates with 1 μL of sterile saline at 200nL per minute. Mice were allowed to heal for two weeks. Mice were then implanted with either placebo or isradipine slow-release pellets (Innovative Research of America). Pellets were designed to release 3μg/g/day of isradipine. Briefly, mice were anaesthetized with isoflurane and a placebo or isradipine pellet was implanted subcutaneously and the wound was closed with sutures. Administration of either vehicle or CNO water was initiated the same day as pellet implantation. Blood was drawn from anaesthetized mice and incubated in tubes containing K3 EDTA solution on ice. Tubes were centrifuged at 4°C at 10,000 *g* for 10 minutes. Collected supernatant was stored at –80°C until analyzed. Samples were sent to Charles River Laboratories for LC-MS/MS bioanalysis.

### Monoamine Quantification

Striatal tissues flash frozen for Visium Spatial Transcriptomics (described below) were used for monoamine quantification. Briefly, flash frozen brains were embedded in OCT and mounted on a cryostat to collect dorsal striatal punches. Punches were stored at −80°C before shipment to the Vanderbilt Neurochemistry Core for catecholamine quantification by HPLC (*27*).

### Immunohistochemistry

Animals were anesthetized with 2,2,2-tribromoethanol and perfused with PBS followed by 4% paraformaldehyde (PFA) in PBS. Intact brains were removed, post-fixed in 4% PFA overnight at 4°C and cryoprotected in 30% sucrose. 40-μm-thick coronal sections were cut on a sliding microtome (Leica) and stored in cryoprotectant (30% Ethylene Glycol (Sigma), 30% Glycerol (Fisher scientific) in PBS).

A detailed protocol for immunofluorescence can be found at dx.doi.org/10.17504/protocols.io.kxygx38owg8j/v1. Brain sections were rinsed with PBS followed by 0.2% Triton X-100 in PBS. The sections were then transferred to a blocking solution containing 4% fetal bovine serum (JR Scientific) and 0.2% Triton X-100 in PBS for 1 hour. Following overnight incubation in primary antibody, sections were rinsed in 0.2% Triton X-100 in PBS and incubated for 2 hours with a suitable secondary antibody. Sections were rinsed again in 0.2% Triton X-100 before mounting and coverslipping with antifade mounting medium (Vector Laboratories H1400, H1500).

A detailed protocol for peroxidase staining can be found at dx.doi.org/10.17504/protocols.io.n92ldm127l5b/v1. Sections were quenched with 3% H_2_O_2_ and 10% methanol in PBS and blocked in 10% fetal bovine calf serum, 3% BSA and 1% glycine in PBS with 0.5% Triton X-100. They were incubated with primary antibody followed by biotinylated secondary and subsequently streptavidin-conjugated horseradish peroxidase (HRP) (1:500; Vectastain ABC kit, Vector Laboratories). Immunostaining was visualized with hydrogen peroxide and 1 3,3′-diaminobenzidine (DAB, Sigma).

The following primary antibodies were used: rabbit anti-DsRed (1:1000; Takara, RRID: AB_3083500), rabbit anti-tyrosine hydroxylase (1:1000; AB152, Millipore, RRID: AB_390204), mouse anti-NeuN (Millipore MAB 377, RRID: AB_2298772), and chicken anti-mCherry (Abcam ab205402, RRID: AB_2722769). Secondary antibodies Alexafluor goat anti-mouse 647 (Thermo Fisher Scientific Cat# A-21235, RRID:AB_2535804), anti-rabbit 488 (Thermo Fisher Scientific Cat# A11034, RRID:AB_2576217), anti-rabbit 594 (Thermo Fisher Scientific Cat# A11037, RRID:AB_2534095), or anti-chicken 594 (Thermo Fisher Scientific Cat# A-11042, RRID:AB_2534099) IgG were used (1:500). A biotinylated goat anti-rabbit IgG (1:300, Vector Laboratories, BA-1000, RRID:AB_2313606) was used for peroxidase staining.

Stained samples were visualized using an automated fluorescence microscope (Keyence BZ7000) and a laser-scanning confocal microscope (Zeiss LSM880).

### Stereology

Total numbers of TH- and mCherry-positive neurons were quantified blinded to groups. Region selection of SN and VTA was done under 5× magnification following the definition by Fu and colleagues (*97*). Imaging was done under 63× magnification by a Zeiss Imager microscope (Carl Zeiss Axio Imager A1) equipped with an XYZ computer-controlled motorized stage and an EOS rebel T5i Digital Camera (Canon), and counting was done using MBF Bioscience’s Stereo Investigator (RRID: SCR_024705). Counting frame size was 60 × 60 μm and systematic random sampling (SRS) grid was 130 × 130 μm, with a section interval of 6.

### Optical Density Analysis

A detailed protocol for optical density analysis can be found at dx.doi.org/10.17504/protocols.io.81wgbxo2nlpk/v1. Images of DAB-stained striatal sections were taken with an automated light microscope (Keyence) by a blinded experimenter. ImageJ (RRID: SCR_002285) was calibrated for optical density and subsequently used to draw ROIs around striatal areas and to measure mean optical density. The Allen Brain Atlas was used as a reference brain atlas (Allen Mouse Brain Atlas, mouse.brain-map.org and atlas.brain-map.org). Optical density values were background subtracted using an adjacent brain region with low TH expression levels, the piriform cortex.

### Spatial Genomics

#### Mouse Visium Spatial Gene Expression

Spatial transcriptomics were acquired with Visium spatial gene expression kits (10X Genomics). Sample preparation, sample imaging, and library generation were completed in accordance with 10X Spatial Gene Expression protocols and as previously published (*98*). Briefly, fresh brain tissue was flash frozen in an isopentane bath cooled to −80°C using dry ice. The brain tissue was then embedded in Optimal Cutting Temperature (OCT) compound (Tissue-Tek 62550-12). A cryostat was used to obtain a 10 µm thin section from the midbrain that was then mounted onto a 10X spatial gene expression slide. Sections were stained with TH, NeuN, and DsRed to visualize mCherry (striatal sections only), and Hoechst 33342 before imaging on a Leica Aperio Versa slide scanner. The cDNA libraries were generated at the Gladstone Genomics Core. Libraries were sequenced at the UCSF Center for Advanced Technology on an Illumina NovaSeq 600 on an SP flow cell. Alignment of the sequencing data with spatial data from the Visium slide was completed with the 10X Space Ranger software (10x Genomics, RRID: SCR_023571). The 10X Loupe Browser software (RRID: SCR_018555) was then used to identify six anatomical regions of interest: (1) SN, (2) VTA, and (3) thalamus in the midbrain, and (4) CP, (5) LSX, and (6) white matter tracts in the striatum. RNA capture areas corresponding to each anatomical region were selected for analysis based on their spatial proximity to the anatomical regions and on the expression of known genetic markers. SN and VTA genetic markers included TH, DAT, and VMAT2; thalamus markers included Prkcd, Ptpn3, and Synpo2; CP was identified with DARPP32; the LSX was identified with PRKCD; white matter was identified with MBP. Demarcation of SN and VTA was done according to Fu and colleagues (*97*). Capture areas that expressed high levels of astrocyte markers (Gfap, Aldh1l1) and microglia markers (Aif1, P2ry12) were excluded. SN and VTA capture areas also had to contain at least one complete DA neuron soma. Roughly 5-10 neuronal cell bodies fit into a single capture area for SN and VTA with 2-49 capture areas per VTA, 1-7 capture areas per SN, 0-39 capture areas per thalamus, 357-560 capture areas per CP, 6-93 capture areas per LSX, and 21-72 capture areas per white matter tracts. Gene expression levels were exported to GraphPad Prism and Microsoft Excel (Microsoft, RRID: SCR_016137). Gene rankings for hit analysis were established using the fold change score (FCS) and signal-to-noise score (SNS). Equations for these scores are given as:

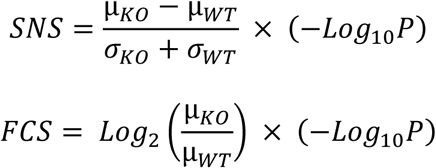

where µ is the average gene expression, σ is the standard deviation, and P is the p-value derived from a t-test. Genes with a p-value < 0.05 were highlighted as differentially expressed genes of interest. The expression levels for these genes of interest were probed in SN, VTA, and the thalamus to identify a subset of genes with DA region-specific changes. Pathway analysis was done on all hits that appeared in both scoring metrics using the Enrichr webtool (RRID: SCR_001575) (*99–101*). Code for Visium spatial transcriptomics analysis can be found on GitHub at https://github.com/yoshitakasei/NakamuraVisiumR-Analysis.

#### Principal Components Analysis

The R function prcomp was used for principal component analysis, with the median normalized gene expression level of each gene as the input. Only genes that were expressed in all regions (for SN and VTA) and whose expression was >0 were included.

#### Neuronal Activity Predictions

To quantify neuronal activity, we utilized NEUROeSTIMator (*47*), a deep learning model that generates a singular activity score following whole transcriptome reconstruction of 22 neuronal activity markers. This score has been shown to correlate significantly with electrophysiological features of hyper-excitability. We applied the NEUROeSTIMator algorithm to the Visium count matrix (species = “mmusculus”). Visium spatial transcriptomic capture areas within the SN and VTA were defined as described above, and neighboring spots were included using the RegionNeighbors (mode = “all_inner_outer”) function of semla R statistical program (*102*). The predicted activity scores were normalized by division of the mean activity scores of the total tissue section they were derived. Group-wise normalized predicted activity scores were compared using the Kolmogorov-Smirnov test (KS-test).

#### Mouse Xenium Spatial Gene Expression

The 10X Genomics Xenium In Situ single cell spatial imaging platform was completed with Stanford Genomics in accordance with the 10X Xenium In Situ protocol (10X Genomics CG000581, CG000582, and CG000584 (*103*)). Flash-frozen tissues embedded in OCT were mounted to Xenium Slides (10X Genomics 1000460) and imaged on the 10X Genomics Xenium Analyzer instrument (RRID:SCR_023910). The Xenium Analyzer performs Xenium Onboard Analysis (RRID:SCR_026158), which includes image pre-processing, puncta detection, transcript decoding, and quality score assignment. Transcript counts were restricted to the cell nucleus using the 10X Genomics Xenium Ranger v2.0.1 software suite.

A post-hoc immunofluorescence stain for mCherry was performed according to 10X Genomics protocols (10X Genomics CG000709) using the same DsRed primary and Alexa-594 secondary mentioned above. Images acquired at 40X using a laser-scanning confocal microscope (Zeiss LSM 880) were converted to pyramidal OME-TIFF files with QuPath v0.5.1 (RRID:SCR_018257). Images were aligned in the Xenium Explorer v3.2.0 (RRID:SCR_025847) according to 10X Genomics Image Alignment protocol adapted from Horn’s absolute orientation algorithm (*104*).

Nuclei of dopaminergic neurons were identified via expression of TH, DAT, or VMAT2. Of the dopaminergic nuclei identified, only the nuclei surrounded by DsRed fluorescence (indicative of successful viral infection) were selected for transcript count analysis. Transcript counts for dopaminergic neurons infected with virus were extracted and exported using Seurat v5.1.0. The exported average transcript counts per nucleus were visualized in GraphPad Prism. Code for Xenium spatial transcriptomics analysis can be found on GitHub at https://github.com/yoshitakasei/Pilot_Xenium_w_mCherry_Marker.git.

#### Human Spatial Gene Expression

Cohort materials: Formalin-fixed paraffin-embedded (FFPE) midbrain sections were collected from individuals with pathologically verified early-stage Parkinson’s Disease (ePD) (n=10) and individuals without any neurological or neuropathological conditions (n=10) through the New South Wales (NSW) Brain Banks, as detailed in Supplementary Table 2. The research protocol received ethical clearance from the University of Sydney Human Research Ethics Committee (approval number 2021/845). All PD subjects exhibited a positive response to levodopa and met the UK Brain Bank Clinical Criteria for PD diagnosis (*105*) without any other neurodegenerative disorders. Control subjects demonstrated no signs of Lewy body pathology, and ePD subjects had Braak stage IV Lewy body pathology, in accordance with established criteria (*106, 107*).

Sequencing and data processing: FFPE sections were stained using TH antibody (BioLegend, cat# 818008, 1:50 RRID: AB_2801155) and processed using the Nanostring GeoMx® Digital Spatial Profiler (RRID: SCR_021660), as per manufacturer instructions, to obtain TH+ masked transcripts. Sequencing libraries for the whole transcriptome were constructed using the Human Whole Transcriptome Atlas (GeoMx Hu WTA) following manufacturer instructions. Technical replicates were consolidated using the Linux ‘cat’ command, and alignment and feature counting were performed using the GeoMx NGS analysis pipeline (version 2.0.21) executed on the Illumina BaseSpace platform (Illumina, RRID: SCR_011881). Quality control was implemented using R statistical programming language with the cutoffs; minSegmentReads: 1000, percentTrimmed: 80%, percentStitched: 80%, percentAligned: 75%, percentSaturation: 50%, minNegativeCount: 1, maxNTCCount: 10,000, minNuclei: 20, minArea: 1000. The minimum gene detection rate across all samples was set at 1%. The minimum gene detection rate per sample was set to 1%. Recently, Van Hijfte and colleagues reviewed the recommended Nanostring GeoMx Q3 normalization technique (*108*), observing that Q3 normalization failed to correct for large differences in the signal (gene expression) to noise (neg probes) observed between samples and recommended quantile normalization. We performed preliminary experiments of 5 normalization techniques; Q3 normalization, background normalization, quantile normalization, SCTransform and normalizing for total library size. We observed that quantile normalization displayed the lowest absolute MA plot correlation and least significance following Kolmogorov Smirnov test. Hence, quantile normalization was implemented, and batches (slides) were corrected using Harmony (*109*), DEGs identified by spatial transcriptomic analysis of mouse model were evaluated between control and ePD TH+ masked Regions of Interest (ROIs) from the SNc ventral tier. The congruence between directional changes of mouse and human was quantified using a chi-square test. To assess differential gene expression between control (CTR) and Parkinson’s disease (PD) samples, we employed the ‘limma voom’ methodology, incorporating covariates diagnosis, age, post-mortem delay (PMD), RNA integrity number equivalent (DV200), sex, DSP processing plate, and brain bank ID (*110*). This analysis was conducted using the R statistical software environment with all scripts are available at https://github.com/zchatt/ASAP-SpatialTranscriptomics/tree/Rademacher_2024 and processed and raw datasets are available at <u>10.5281/zenodo.10499186</u>.

### Quantification and Statistical Analysis

All statistical analyses including the n, what n represents, description of error bars, statistical tests used and level of significance are found in the figure legends. Two-way repeated-measures ANOVA followed by Holm-Sidak multiple comparisons was used for comparing vehicle vs CNO groups. One-way ANOVA followed by Holm-Sidak multiple comparisons was used for comparing multiple groups in mouse spatial transcriptomic data. Three-way ANOVA followed by Holm-Sidak multiple comparisons was used for assessing chronic isradipine histology. T-test or permutation (non-parametric) analysis was used for ex vivo electrophysiology data. Mouse differentially expressed genes were ranked according to the defined SNS and FCS metrics. Congruence between directional changes of mouse and human was quantified using a chi-square test. ‘Limma voom’ methodology was used to assess differential gene expression between control and PD samples, incorporating covariates (see Human Spatial Gen Expression). All analyses were performed using GraphPad Prism version 9 (RRID: SCR_002798), Microsoft Excel, and R version 4.2.2 (RRID: SCR_000432).

## Data, Materials, and Software Availability

Datasets are available at https://doi.org/10.5281/zenodo.10499027. All other data are included in the manuscript and/or supporting information.

## Acknowledgements

We acknowledge Haru Yamamoto and James Maas for assistance preparing the manuscript, and Kathryn Claiborn for helping edit the manuscript and Erica Delin for administrative assistance. We thank Saheli Singh for assistance with mouse spatial transcriptomics analysis, Robert Edwards, Zayd Khaliq, Talia Lerner and Christopher Ford from “Team Edwards” for feedback on experiments, and Yutau Liu for technical assistance.

This research was funded in whole or in part by Aligning Science Across Parkinson’s (ASAP-020529) through the Michael J. Fox Foundation for Parkinson’s Research (MJFF). For the purpose of open access, the author has applied a CC BY public copyright license to all Author Accepted Manuscripts arising from this submission. This work was also supported by NIH (RO1NS091902 and RF1AG064170, KN; R01DA030529, EBM; F31NS137765, KR), the State of California for medical research on alcohol and substance abuse through the State of California, the Joan and David Traitel Family Trust and Betty Brown’s Family, a Burroughs-Wellcome Fund Award (KN), and the Gladstone Institutes. This work was also supported by the Hillblom Foundation (ZD) and a Berkelhammer Award for Excellence in Neuroscience (ZD).

## References

1. J. R. G. Hollerman, A. A., The effects of dopamine-depleting brain lesions on the electrophysiological activity of rat substantia nigra dopamine neurons. Brain Research 533, 203–212 (1990).

2. M. J. Zigmond, A. L. Acheson, M. K. Stachowiak, E. M. Stricker, Neurochemical compensation after nigrostriatal bundle injury in an animal model of preclinical parkinsonism. Arch Neurol 41, 856–861 (1984).

3. Y. Agid, F. Javoy, J. Glowinski, Hyperactivity of remaining dopaminergic neurones after partial destruction of the nigro-striatal dopaminergic system in the rat. Nat New Biol 245, 150–151 (1973).

4. F. Hefti, E. Melamed, R. J. Wurtman, Partial lesions of the dopaminergic nigrostriatal system in rat brain: biochemical characterization. Brain Res 195, 123–137 (1980).

5. W. Q. Zhang et al., Increased dopamine release from striata of rats after unilateral nigrostriatal bundle damage. Brain Res 461, 335–342 (1988).

6. L. Chen et al., Chronic, systemic treatment with a metabotropic glutamate receptor 5 antagonist in 6-hydroxydopamine partially lesioned rats reverses abnormal firing of dopaminergic neurons. Brain Res 1286, 192–200 (2009).

7. M. K. K. J. R. W. S. Stachowiak, E. M.; Zigmond, M. J., Increased dopamine efflux from striatal slices during development and after nigrostriatal bundle damage. J Neurosci 7, 1648–1654 (1987).

8. P. Gonzalez-Rodriguez et al., Disruption of mitochondrial complex I induces progressive parkinsonism. Nature 599, 650–656 (2021).

9. M. W. Bishop et al., Hyperexcitable substantia nigra dopamine neurons in PINK1- and HtrA2/Omi-deficient mice. J Neurophysiol 104, 3009–3020 (2010).

10. J. Schiemann et al., K-ATP channels in dopamine substantia nigra neurons control bursting and novelty-induced exploration. Nat Neurosci 15, 1272–1280 (2012).

11. H. Morikawa, C. A. Paladini, Dynamic regulation of midbrain dopamine neuron activity: intrinsic, synaptic, and plasticity mechanisms. Neuroscience 198, 95–111 (2011).

12. C. H. Good et al., Impaired nigrostriatal function precedes behavioral deficits in a genetic mitochondrial model of Parkinson’s disease. FASEB J 25, 1333–1344 (2011).

13. M. Lin et al., In Parkinson’s patient-derived dopamine neurons, the triplication of alpha-synuclein locus induces distinctive firing pattern by impeding D2 receptor autoinhibition. Acta Neuropathol Commun 9, 107 (2021).

14. M. Regoni et al., Pharmacological antagonism of kainate receptor rescues dysfunction and loss of dopamine neurons in a mouse model of human parkin-induced toxicity. Cell Death Dis 11, 963 (2020).

15. A. Dagra et al., alpha-Synuclein-induced dysregulation of neuronal activity contributes to murine dopamine neuron vulnerability. NPJ Parkinsons Dis 7, 76 (2021).

16. S. Janezic et al., Deficits in dopaminergic transmission precede neuron loss and dysfunction in a new Parkinson model. Proc Natl Acad Sci U S A 110, E4016–4025 (2013).

17. P. D. Dodson et al., Representation of spontaneous movement by dopaminergic neurons is cell-type selective and disrupted in parkinsonism. Proc Natl Acad Sci U S A 113, E2180–2188 (2016).

18. M. Subramaniam et al., Mutant alpha-synuclein enhances firing frequencies in dopamine substantia nigra neurons by oxidative impairment of A-type potassium channels. J Neurosci 34, 13586–13599 (2014).

19. J. S. Chou et al., (G2019S) LRRK2 causes early-phase dysfunction of SNpc dopaminergic neurons and impairment of corticostriatal long-term depression in the PD transgenic mouse. Neurobiol Dis 68, 190–199 (2014).

20. V. M. Nemani et al., Increased expression of alpha-synuclein reduces neurotransmitter release by inhibiting synaptic vesicle reclustering after endocytosis. Neuron 65, 66–79 (2010).

21. H. Bergman, T. Wichmann, B. Karmon, M. R. DeLong, The primate subthalamic nucleus. II. Neuronal activity in the MPTP model of parkinsonism. J Neurophysiol 72, 507–520 (1994).

22. H. Braak, K. Del Tredici, Poor and protracted myelination as a contributory factor to neurodegenerative disorders. Neurobiol Aging 25, 19–23 (2004).

23. P. Tagliaferro, R. E. Burke, Retrograde Axonal Degeneration in Parkinson Disease. J Parkinsons Dis 6, 1–15 (2016).

24. D. Haddad, K. Nakamura, Understanding the susceptibility of dopamine neurons to mitochondrial stressors in Parkinson’s disease. FEBS Lett 589, 3702–3713 (2015).

25. R. Betarbet et al., Chronic systemic pesticide exposure reproduces features of Parkinson’s disease. Nat Neurosci 3, 1301–1306 (2000).

26. Z. Doric, K. Nakamura, Mice with disrupted mitochondria used to model Parkinson’s disease. Nature 599, 558–560 (2021).

27. A. Berthet et al., Loss of mitochondrial fission depletes axonal mitochondria in midbrain dopamine neurons. J Neurosci 34, 14304–14317 (2014).

28. H. Li et al., Longitudinal tracking of neuronal mitochondria delineates PINK1/Parkin-dependent mechanisms of mitochondrial recycling and degradation. Sci Adv 7, (2021).

29. D. C. German, K. F. Manaye, P. K. Sonsalla, B. A. Brooks, Midbrain dopaminergic cell loss in Parkinson’s disease and MPTP-induced parkinsonism: sparing of calbindin-D28k-containing cells. Ann N Y Acad Sci 648, 42–62 (1992).

30. S. J. Kish, K. Shannak, O. Hornykiewicz, Uneven pattern of dopamine loss in the striatum of patients with idiopathic Parkinson’s disease. Pathophysiologic and clinical implications. N Engl J Med 318, 876–880 (1988).

31. Z. M. Khaliq, B. P. Bean, Pacemaking in dopaminergic ventral tegmental area neurons: depolarizing drive from background and voltage-dependent sodium conductances. J Neurosci 30, 7401–7413 (2010).

32. W. Matsuda et al., Single nigrostriatal dopaminergic neurons form widely spread and highly dense axonal arborizations in the neostriatum. J Neurosci 29, 444–453 (2009).

33. D. J. Surmeier, P. T. Schumacker, Calcium, bioenergetics, and neuronal vulnerability in Parkinson’s disease. J Biol Chem 288, 10736–10741 (2013).

34. B. Piallat, A. Benazzouz, A. L. Benabid, Subthalamic nucleus lesion in rats prevents dopaminergic nigral neuron degeneration after striatal 6-OHDA injection: behavioural and immunohistochemical studies. Eur J Neurosci 8, 1408–1414 (1996).

35. H. Bergman, T. Wichmann, M. R. DeLong, Reversal of experimental parkinsonism by lesions of the subthalamic nucleus. Science 249, 1436–1438 (1990).

36. K. A. B. Zaghloul, J. A.; Weidemann, C. T.; McGill, K.; Jaggi, J. L.; Baltuch, G. H.; Kahana, M. J., Human Substantia Nigra Neurons Encode Unexpected Financial Rewards. Science 323, 1496–1499 (2009).

37. Q. M. Wang, Y. Y. Xu, S. Liu, Z. G. Ma, Isradipine attenuates MPTP-induced dopamine neuron degeneration by inhibiting up-regulation of L-type calcium channels and iron accumulation in the substantia nigra of mice. Oncotarget 8, 47284–47295 (2017).

38. E. Ilijic, J. N. Guzman, D. J. Surmeier, The L-type channel antagonist isradipine is neuroprotective in a mouse model of Parkinson’s disease. Neurobiol Dis 43, 364–371 (2011).

39. M. Verma, B. N. Lizama, C. T. Chu, Excitotoxicity, calcium and mitochondria: a triad in synaptic neurodegeneration. Transl Neurodegener 11, 3 (2022).

40. J. A. da Silva, F. Tecuapetla, V. Paixao, R. M. Costa, Dopamine neuron activity before action initiation gates and invigorates future movements. Nature 554, 244–248 (2018).

41. X. Jin, R. M. Costa, Start/stop signals emerge in nigrostriatal circuits during sequence learning. Nature 466, 457–462 (2010).

42. A. V. Kravitz et al., Regulation of parkinsonian motor behaviours by optogenetic control of basal ganglia circuitry. Nature 466, 622–626 (2010).

43. P. A. Kumar et al., Chemogenetic Attenuation of Acute Nociceptive Signaling Enhances Functional Outcomes Following Spinal Cord Injury. J Neurotrauma 41, 1060–1076 (2024).

44. S. Wang, Y. Tan, J. E. Zhang, M. Luo, Pharmacogenetic activation of midbrain dopaminergic neurons induces hyperactivity. Neurosci Bull 29, 517–524 (2013).

45. J. H. Kordower et al., Disease duration and the integrity of the nigrostriatal system in Parkinson’s disease. Brain 136, 2419–2431 (2013).

46. B. L. Roth, DREADDs for Neuroscientists. Neuron 89, 683–694 (2016).

47. E. Bahl et al., Using deep learning to quantify neuronal activation from single-cell and spatial transcriptomic data. Nat Commun 15, 779 (2024).

48. R. Donato, Intracellular and extracellular roles of S100 proteins. Microsc Res Tech 60, 540–551 (2003).

49. F. Lang, N. Strutz-Seebohm, G. Seebohm, U. E. Lang, Significance of SGK1 in the regulation of neuronal function. J Physiol 588, 3349–3354 (2010).

50. A. Filipek, W. Lesniak, S100A6 and Its Brain Ligands in Neurodegenerative Disorders. Int J Mol Sci 21, (2020).

51. S. Iwata, M. Nomoto, H. Morioka, A. Miyata, Gene expression profiling in the midbrain of striatal 6-hydroxydopamine-injected mice. Synapse 51, 279–286 (2004).

52. C. C. Stichel et al., sgk1, a member of an RNA cluster associated with cell death in a model of Parkinson’s disease. Eur J Neurosci 21, 301–316 (2005).

53. A. R. Means, M. F. VanBerkum, I. Bagchi, K. P. Lu, C. D. Rasmussen, Regulatory functions of calmodulin. Pharmacol Ther 50, 255–270 (1991).

54. Y. J. Hsu, J. G. Hoenderop, R. J. Bindels, TRP channels in kidney disease. Biochim Biophys Acta 1772, 928–936 (2007).

55. J. Xu, Z. P. Pang, O. H. Shin, T. C. Sudhof, Synaptotagmin-1 functions as a Ca2+ sensor for spontaneous release. Nat Neurosci 12, 759–766 (2009).

56. A. Kadkova, J. Radecke, J. B. Sorensen, The SNAP-25 Protein Family. Neuroscience 420, 50–71 (2019).

57. L. P. Jackson et al., A large-scale conformational change couples membrane recruitment to cargo binding in the AP2 clathrin adaptor complex. Cell 141, 1220–1229 (2010).

58. P. W. Ho et al., In vivo overexpression of synaptogyrin-3 promotes striatal synaptic dopamine uptake in LRRK2(R1441G) mutant mouse model of Parkinson’s disease. Brain Behav 13, e2886 (2023).

59. J. T. Bendor, T. P. Logan, R. H. Edwards, The function of alpha-synuclein. Neuron 79, 1044–1066 (2013).

60. C. S. Chan et al., ’Rejuvenation’ protects neurons in mouse models of Parkinson’s disease. Nature 447, 1081–1086 (2007).

61. E. Dragicevic et al., Cav1.3 channels control D2-autoreceptor responses via NCS-1 in substantia nigra dopamine neurons. Brain 137, 2287–2302 (2014).

62. J. N. Guzman et al., Systemic isradipine treatment diminishes calcium-dependent mitochondrial oxidant stress. J Clin Invest 128, 2266–2280 (2018).

63. E. Carpene, G. Andreani, G. Isani, Metallothionein functions and structural characteristics. J Trace Elem Med Biol 21 **Suppl 1**, 35–39 (2007).

64. R. F. Fernandez, J. M. Ellis, Acyl-CoA synthetases as regulators of brain phospholipid acyl-chain diversity. Prostaglandins Leukot Essent Fatty Acids 161, 102175 (2020).

65. J. S. Dittman, Unc13: a multifunctional synaptic marvel. Curr Opin Neurobiol 57, 17–25 (2019).

66. F. P. Diekstra et al., C9orf72 and UNC13A are shared risk loci for amyotrophic lateral sclerosis and frontotemporal dementia: a genome-wide meta-analysis. Ann Neurol 76, 120–133 (2014).

67. J. P. Quinn, S. E. Kandigian, B. A. Trombetta, S. E. Arnold, B. C. Carlyle, VGF as a biomarker and therapeutic target in neurodegenerative and psychiatric diseases. Brain Commun 3, fcab261 (2021).

68. M. Pereira Luppi et al., Sox6 expression distinguishes dorsally and ventrally biased dopamine neurons in the substantia nigra with distinctive properties and embryonic origins. Cell Rep 37, 109975 (2021).

69. Z. Gaertner, M. Azcorra, D. A. Dombeck, R. Awatramani, Molecular heterogeneity in the substantia nigra: A roadmap for understanding PD motor pathophysiology. Neurobiol Dis 175, 105925 (2022).

70. M. Azcorra et al., Unique functional responses differentially map onto genetic subtypes of dopamine neurons. Nat Neurosci 26, 1762–1774 (2023).

71. D. Galter, S. Buervenich, A. Carmine, M. Anvret, L. Olson, ALDH1 mRNA: presence in human dopamine neurons and decreases in substantia nigra in Parkinson’s disease and in the ventral tegmental area in schizophrenia. Neurobiol Dis 14, 637–647 (2003).

72. A. Eban-Rothschild, G. Rothschild, W. J. Giardino, J. R. Jones, L. de Lecea, VTA dopaminergic neurons regulate ethologically relevant sleep-wake behaviors. Nat Neurosci 19, 1356–1366 (2016).

73. M. Helwig et al., Neuronal hyperactivity-induced oxidant stress promotes in vivo alpha-synuclein brain spreading. Sci Adv 8, eabn0356 (2022).

74. D. G. Harden, A. A. Grace, Activation of dopamine cell firing by repeated L-DOPA administration to dopamine-depleted rats: its potential role in mediating the therapeutic response to L-DOPA treatment. J Neurosci 15, 6157–6166 (1995).

75. B. Liss, R. Bruns, J. Roeper, Alternative sulfonylurea receptor expression defines metabolic sensitivity of K-ATP channels in dopaminergic midbrain neurons. EMBO J 18, 833–846 (1999).

76. G. Bilbao et al., Electrophysiological characterization of substantia nigra dopaminergic neurons in partially lesioned rats: effects of subthalamotomy and levodopa treatment. Brain Res 1084, 175–184 (2006).

77. B. Liss et al., K-ATP channels promote the differential degeneration of dopaminergic midbrain neurons. Nat Neurosci 8, 1742–1751 (2005).

78. A. G. Yee et al., Effects of the Parkinsonian toxin MPP+ on electrophysiological properties of nigral dopaminergic neurons. Neurotoxicology 45, 1–11 (2014).

79. D. R. Curtis, J. W. Phillis, J. C. Watkins, The chemical excitation of spinal neurones by certain acidic amino acids. J Physiol 150, 656–682 (1960).

80. A. A. Grace, B. S. Bunney, Induction of depolarization block in midbrain dopamine neurons by repeated administration of haloperidol: analysis using in vivo intracellular recording. J Pharmacol Exp Ther 238, 1092–1100 (1986).

81. A. A. Grace, B. S. Bunney, H. Moore, C. L. Todd, Dopamine-cell depolarization block as a model for the therapeutic actions of antipsychotic drugs. Trends Neurosci 20, 31–37 (1997).

82. F. J. White, R. Y. Wang, Differential effects of classical and atypical antipsychotic drugs on A9 and A10 dopamine neurons. Science 221, 1054–1057 (1983).

83. J. R. Hollerman, A. A. Grace, Acute haloperidol administration induces depolarization block of nigral dopamine neurons in rats after partial dopamine lesions. Neurosci Lett 96, 82–88 (1989).

84. M. Y. Jing et al., Re-examining the role of ventral tegmental area dopaminergic neurons in motor activity and reinforcement by chemogenetic and optogenetic manipulation in mice. Metab Brain Dis 34, 1421–1430 (2019).

85. N. E. Taylor et al., Optogenetic activation of dopamine neurons in the ventral tegmental area induces reanimation from general anesthesia. Proc Natl Acad Sci U S A 113, 12826–12831 (2016).

86. S. Imai, N. Okayama, M. Shimizu, M. Itoh, Increased intracellular calcium activates serum and glucocorticoid-inducible kinase 1 (SGK1) through a calmodulin-calcium calmodulin dependent kinase kinase pathway in Chinese hamster ovary cells. Life Sci 72, 2199–2209 (2003).

87. K. Szydlowska, M. Tymianski, Calcium, ischemia and excitotoxicity. Cell Calcium 47, 122–129 (2010).

88. R. Sattler, M. P. Charlton, M. Hafner, M. Tymianski, Distinct influx pathways, not calcium load, determine neuronal vulnerability to calcium neurotoxicity. J Neurochem 71, 2349–2364 (1998).

89. A. Abdanipour et al., Neuroprotective effects of selegiline on rat neural stem cells treated with hydrogen peroxide. Biomed Rep 8, 41–46 (2018).

90. M. Elkenani et al., Heat shock protein A4 ablation leads to skeletal muscle myopathy associated with dysregulated autophagy and induced apoptosis. J Transl Med 20, 229 (2022).

91. Y. Yang et al., SIRT1 attenuates neuroinflammation by deacetylating HSPA4 in a mouse model of Parkinson’s disease. Biochim Biophys Acta Mol Basis Dis 1868, 166365 (2022).

92. M. Quik, Smoking, nicotine and Parkinson’s disease. Trends Neurosci 27, 561–568 (2004).

93. J. B. Nourse, Jr., et al., Conserved nicotine-activated neuroprotective pathways involve mitochondrial stress. iScience 24, 102140 (2021).

94. C. Xiao et al., Chronic nicotine selectively enhances alpha4beta2* nicotinic acetylcholine receptors in the nigrostriatal dopamine pathway. J Neurosci 29, 12428–12439 (2009).

95. M. L. Hacker et al., Deep brain stimulation in early-stage Parkinson disease: Five-year outcomes. Neurology 95, e393–e401 (2020).

96. C. M. Backman et al., Characterization of a mouse strain expressing Cre recombinase from the 3’ untranslated region of the dopamine transporter locus. Genesis 44, 383–390 (2006).

97. Y. Fu et al., A cytoarchitectonic and chemoarchitectonic analysis of the dopamine cell groups in the substantia nigra, ventral tegmental area, and retrorubral field in the mouse. Brain Struct Funct 217, 591–612 (2012).

98. Y. J. Sei, M. M. Chaumeil, K. Nakamura, Protocol to combine brain sections from multiple mice into a single block for spatial transcriptomic analyses. STAR Protoc 4, 102617 (2023).

99. E. Y. Chen et al., Enrichr: interactive and collaborative HTML5 gene list enrichment analysis tool. BMC Bioinformatics 14, 128 (2013).

100. M. V. Kuleshov et al., Enrichr: a comprehensive gene set enrichment analysis web server 2016 update. Nucleic Acids Res 44, W90–97 (2016).

101. Z. Xie et al., Gene Set Knowledge Discovery with Enrichr. Curr Protoc 1, e90 (2021).

102. L. Larsson, L. Franzen, P. L. Stahl, J. Lundeberg, Semla: a versatile toolkit for spatially resolved transcriptomics analysis and visualization. Bioinformatics 39, (2023).

103. X. Ma et al., Protocol for Xenium spatial transcriptomics studies using fixed frozen mouse brain sections. STAR Protoc 5, 103420 (2024).

104. B. K. P. Horn, Closed-form solution of absolute orientation using unit quaternions. J. Opt. Soc. Am. A 4, 629–642 (1987).

105. E. Gibbels, [Hitler’s Parkinson syndrome. A posthumous motility analysis of film records of the German Weekly News 1940-1945]. Nervenarzt 59, 521–528 (1988).

106. H. Braak et al., Staging of brain pathology related to sporadic Parkinson’s disease. Neurobiol Aging 24, 197–211 (2003).

107. G. M. Halliday, K. Del Tredici, H. Braak, Critical appraisal of brain pathology staging related to presymptomatic and symptomatic cases of sporadic Parkinson’s disease. J Neural Transm Suppl, 99–103 (2006).

108. L. van Hijfte et al., Alternative normalization and analysis pipeline to address systematic bias in NanoString GeoMx Digital Spatial Profiling data. iScience 26, 105760 (2023).

109. I. Korsunsky et al., Fast, sensitive and accurate integration of single-cell data with Harmony. Nat Methods 16, 1289–1296 (2019).

110. C. W. Law, Y. Chen, W. Shi, G. K. Smyth, voom: Precision weights unlock linear model analysis tools for RNA-seq read counts. Genome Biol 15, R29 (2014).

