## Supplementary Figures and Tables for "Chronic hyperactivation of midbrain dopamine neurons causes preferential dopamine neuron degeneration"

Figure S1

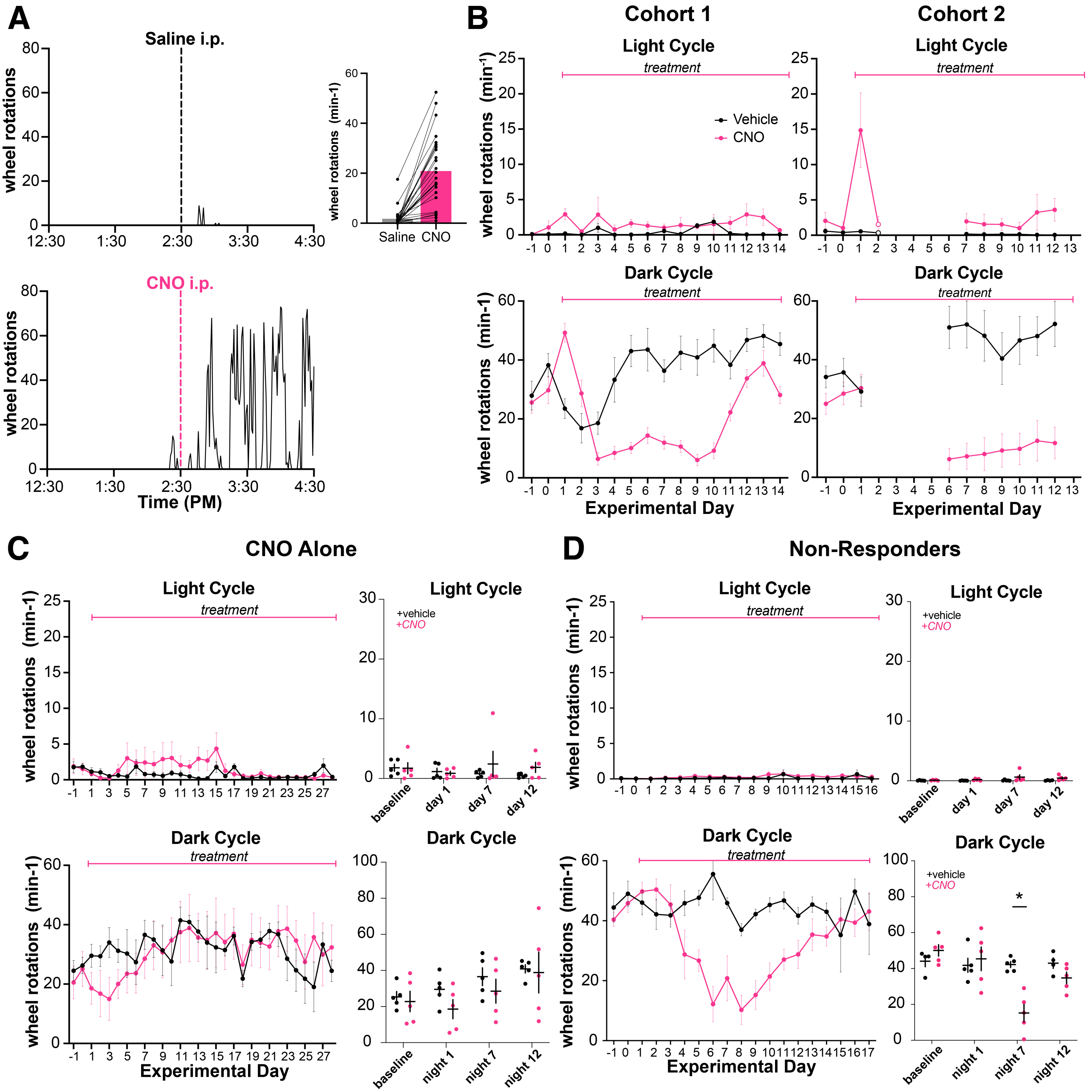
A
Saline i.p.
CNO i.p.
wheel rotations
Time (PM)
12:30
1:30
2:30
3:30
4:30
wheel rotations (min-1)
Saline
CNO
B
Cohort 1
Cohort 2
Light Cycle
Dark Cycle
treatment
Vehicle
CNO
wheel rotations (min-1)
Experimental Day
C
CNO Alone
+vehicle
+CNO
baseline
day 1
day 7
day 12
night 1
night 7
night 12
D
Non-Responders
*

Figure S2

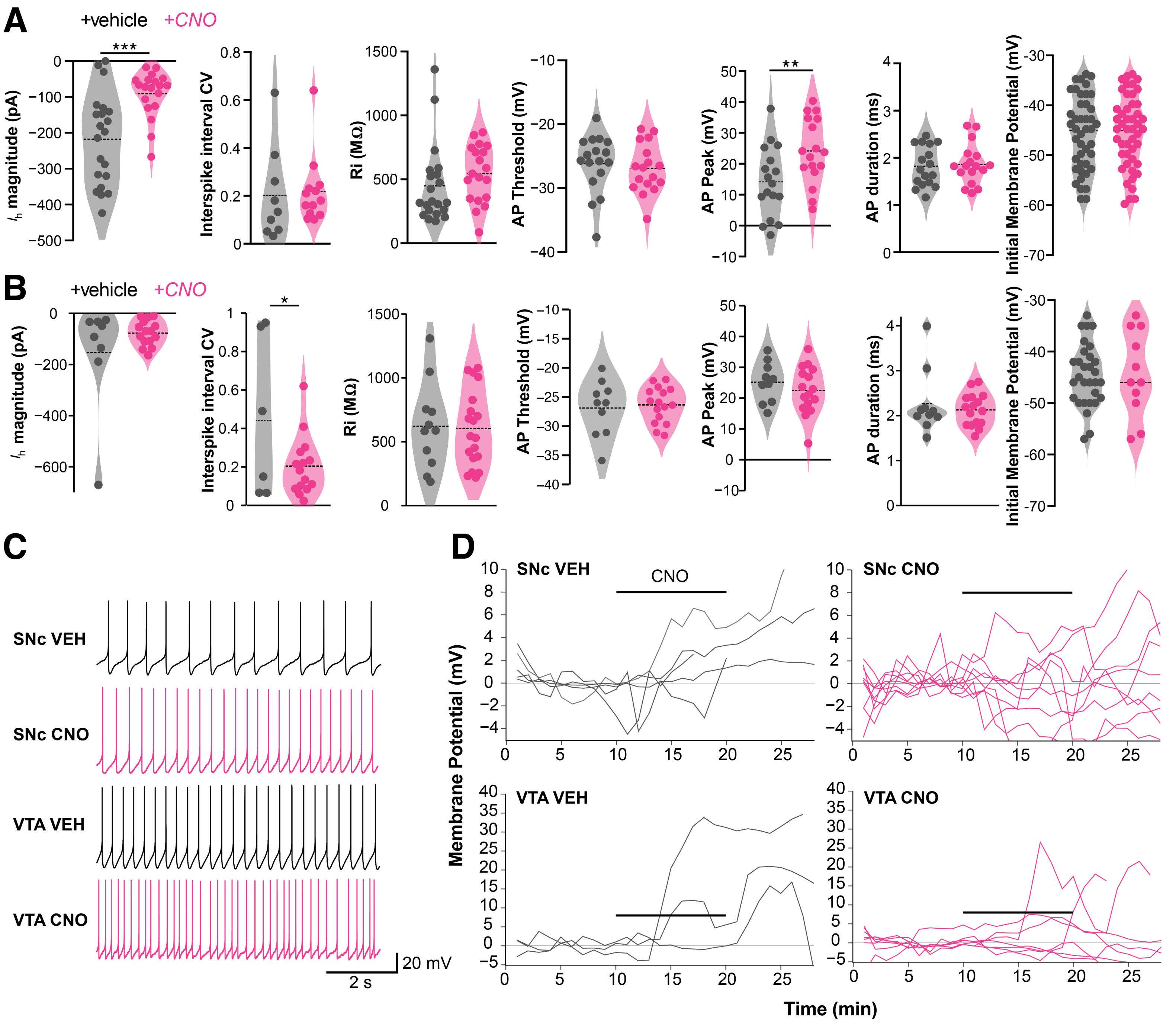
A
+vehicle
+CNO
***
**
$I_h$ magnitude (pA)
Interspike interval CV
Ri (MΩ)
AP Threshold (mV)
AP Peak (mV)
AP duration (ms)
Initial Membrane Potential (mV)
B
+vehicle
+CNO
*
C
SNc VEH
SNc CNO
VTA VEH
VTA CNO
20 mV
2 s
D
SNc VEH
CNO
SNc CNO
VTA VEH
VTA CNO
Membrane Potential (mV)
Time (min)

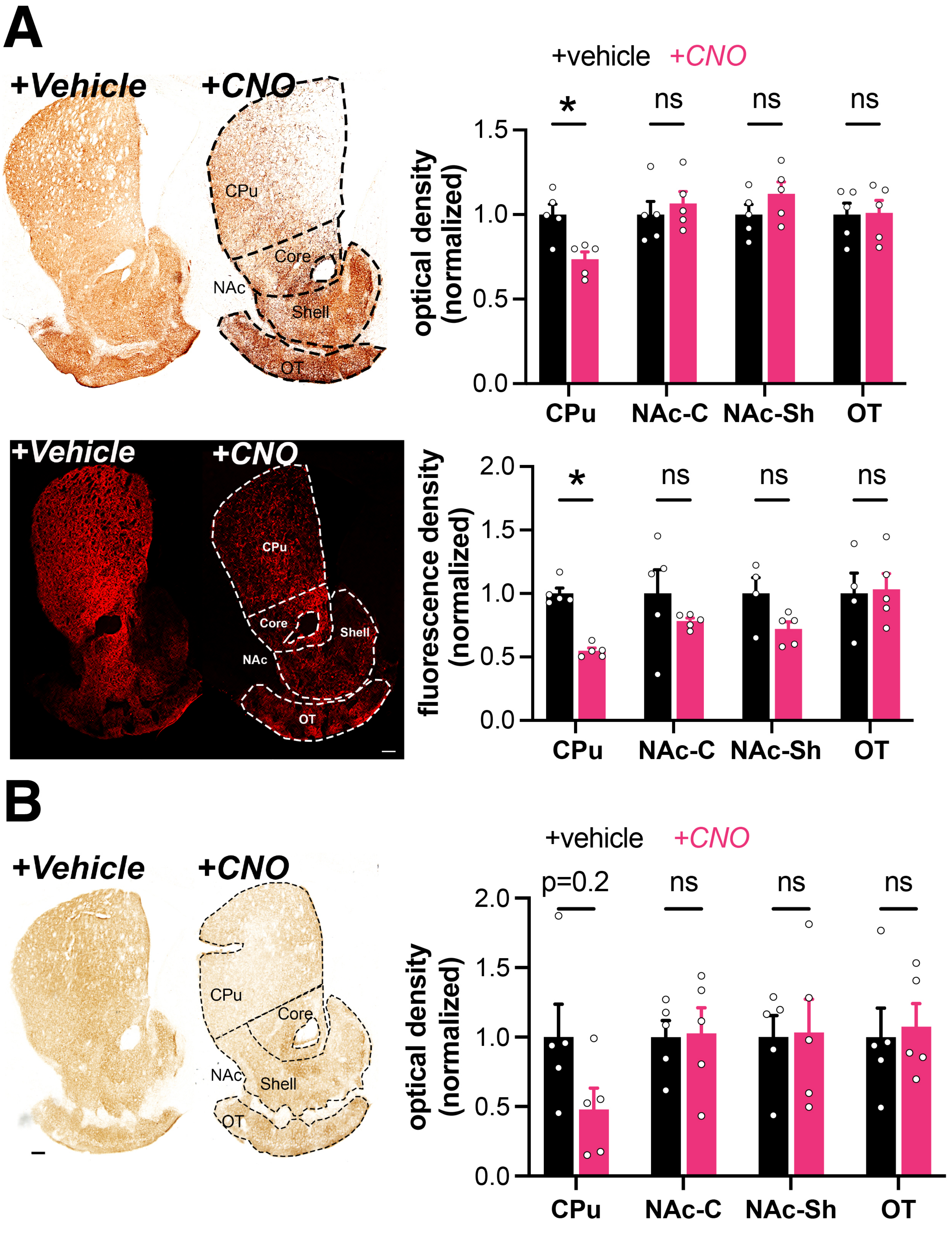
A
+Vehicle
+CNO
CPu
Core
NAc
Shell
OT
+vehicle
+CNO
*
ns
ns
ns
optical density
(normalized)
1.5
1.0
0.5
0.0
CPu
NAc-C
NAc-Sh
OT
+Vehicle
+CNO
CPu
Core
Shell
NAc
OT
*
ns
ns
ns
fluorescence density
(normalized)
2.0
1.5
1.0
0.5
0.0
CPu
NAc-C
NAc-Sh
OT
B
+Vehicle
+CNO
CPu
Core
NAc
Shell
OT
+vehicle
+CNO
p=0.2
ns
ns
ns
optical density
(normalized)
2.0
1.5
1.0
0.5
0.0
CPu
NAc-C
NAc-Sh
OT

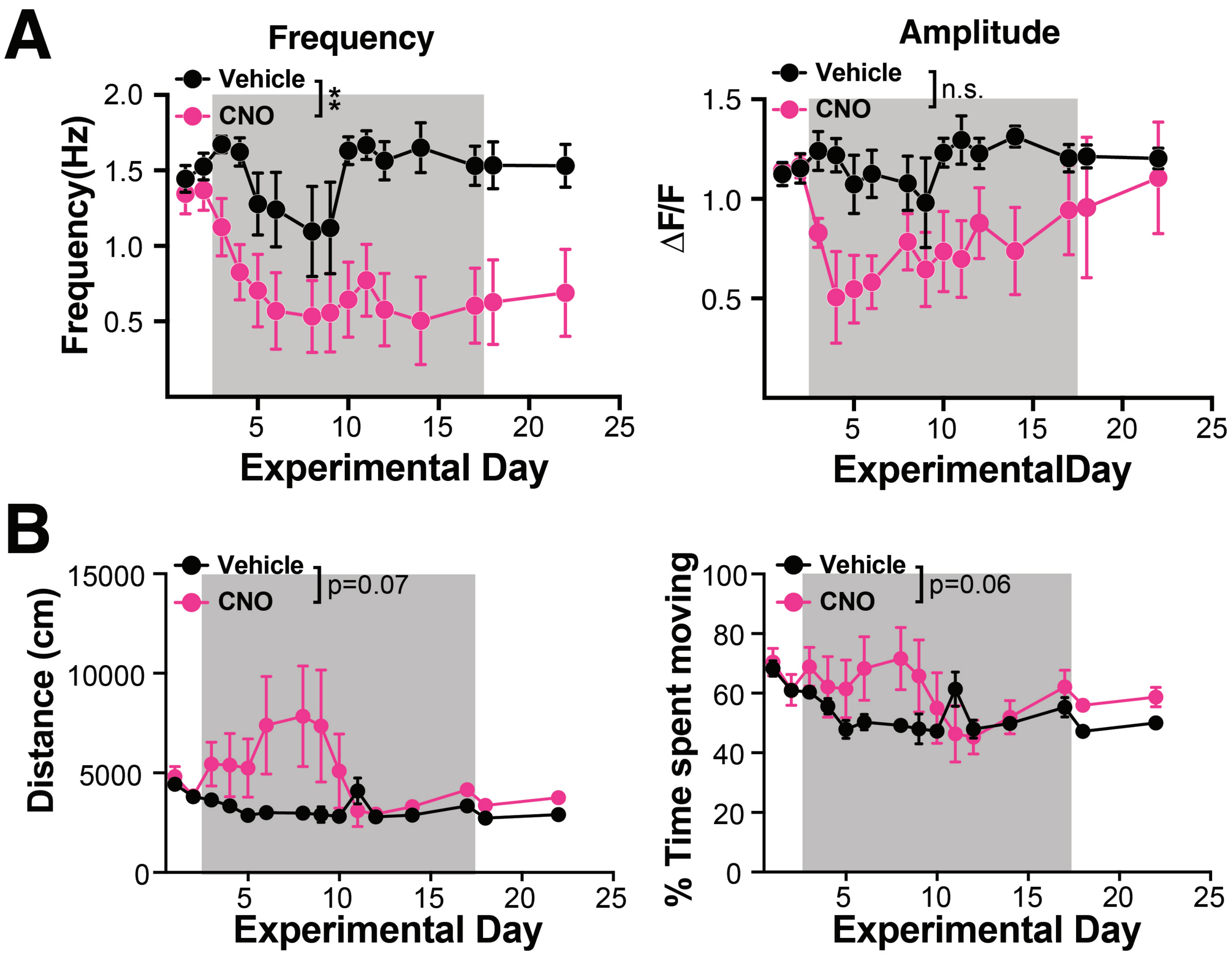
A
Frequency
Vehicle
CNO
**
Frequency(Hz)
2.0
1.5
1.0
0.5
5
10
15
20
25
Experimental Day
Amplitude
Vehicle
CNO
n.s.
ΔF/F
1.5
1.0
0.5
ExperimentalDay
B
Vehicle
CNO
p=0.07
Distance (cm)
15000
10000
5000
0
Experimental Day
Vehicle
CNO
p=0.06
% Time spent moving
100
80
60
40
20
0
Experimental Day

Figure S5

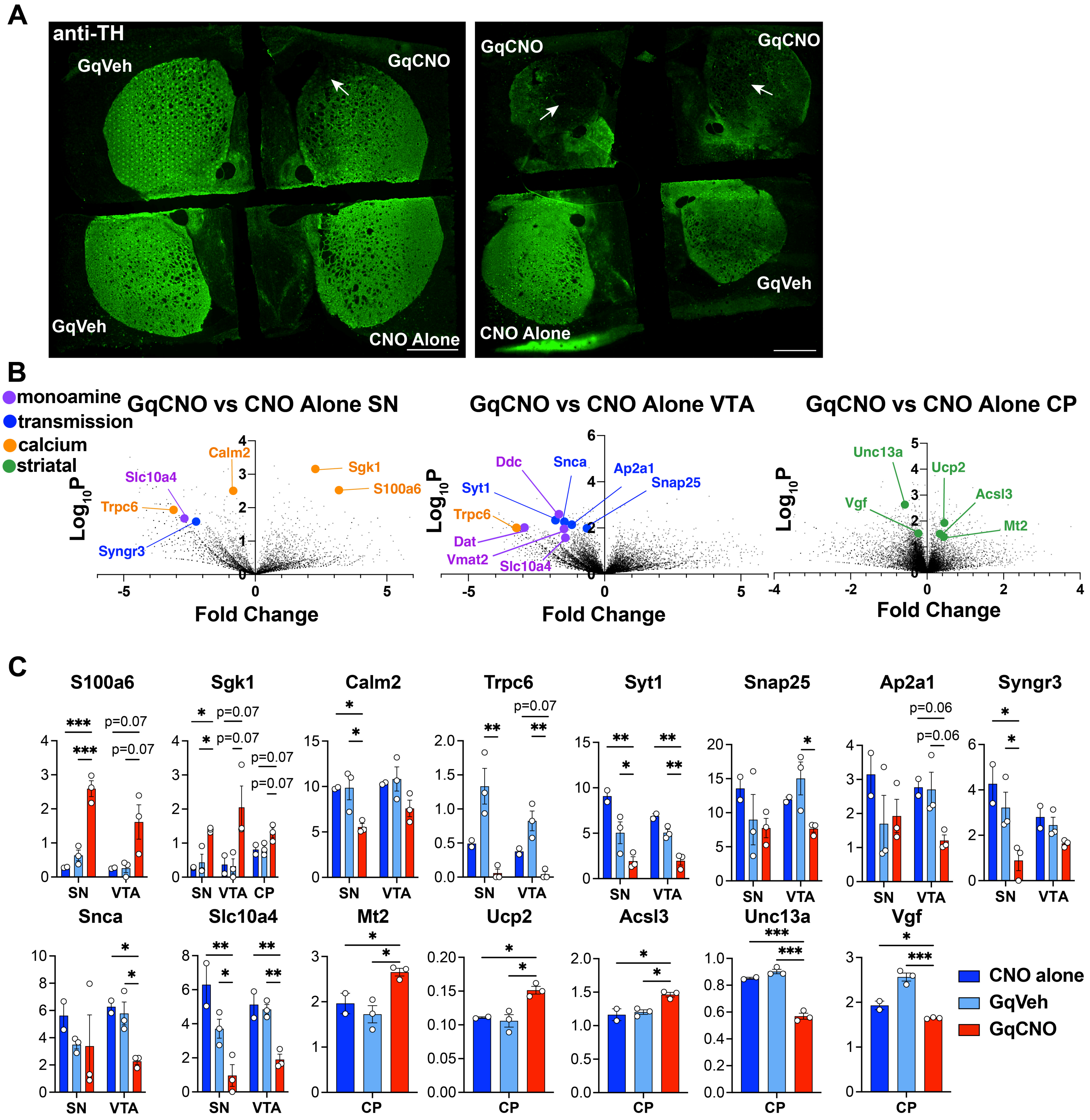
A
anti-TH
GqVeh
GqCNO
GqVeh
CNO Alone
GqCNO
GqCNO
GqVeh
CNO Alone
B
monoamine
transmission
calcium
striatal
GqCNO vs CNO Alone SN
GqCNO vs CNO Alone VTA
GqCNO vs CNO Alone CP
Log10P
Fold Change
Calm2
Sgk1
S100a6
Slc10a4
Trpc6
Syngr3
Ddc
Snca
Ap2a1
Snap25
Syt1
Dat
Vmat2
Slc10a4
Unc13a
Ucp2
Acsl3
Vgf
Mt2
C
S100a6
Sgk1
Calm2
Trpc6
Syt1
Snap25
Ap2a1
Syngr3
Snca
Slc10a4
Mt2
Ucp2
Acsl3
Unc13a
Vgf
p=0.07
p=0.06
SN
VTA
CP
CNO alone
GqVeh
GqCNO

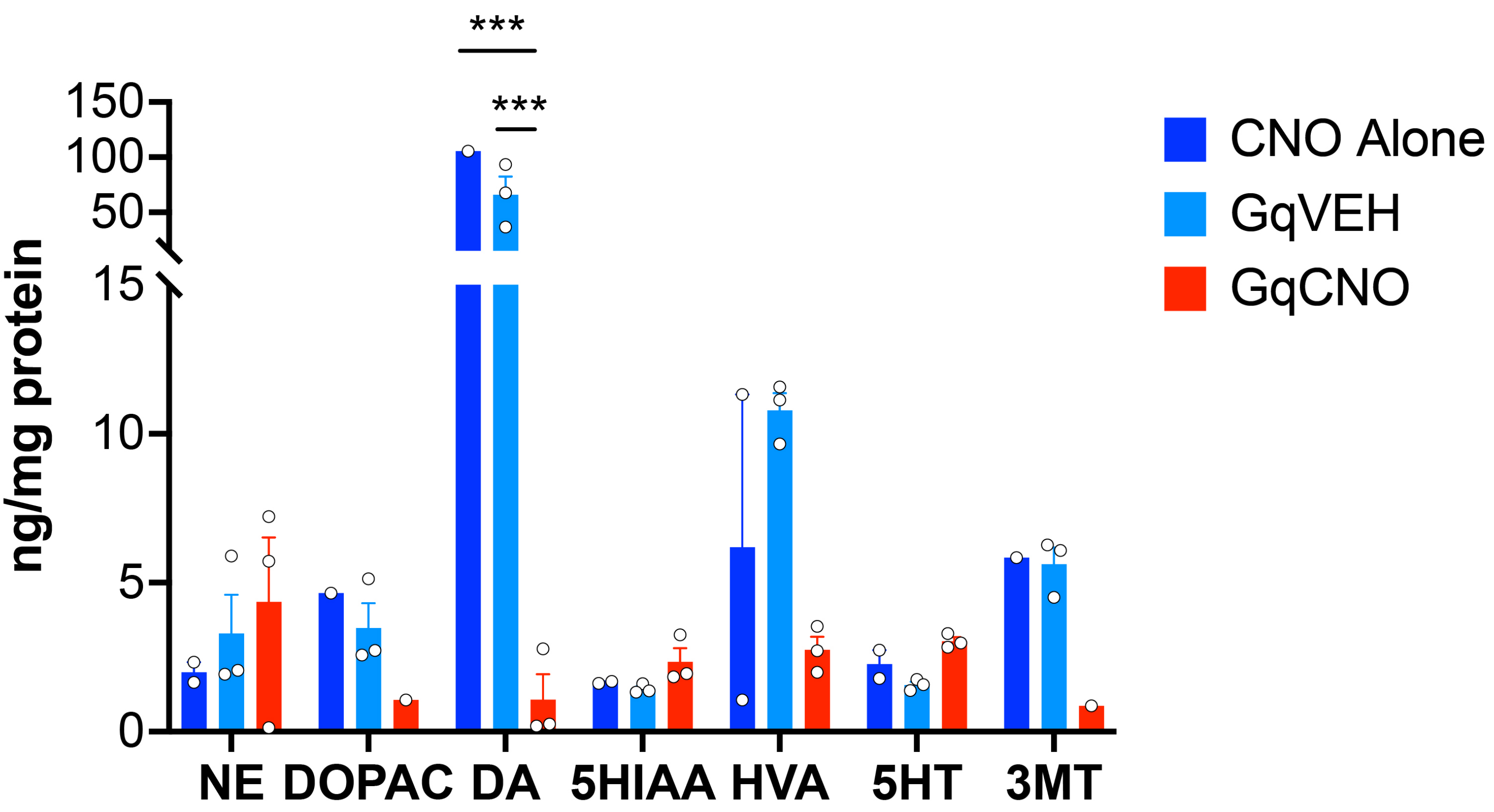
***
***
150
100
50
15
10
5
0
ng/mg protein
NE
DOPAC
DA
5HIAA
HVA
5HT
3MT
CNO Alone
GqVEH
GqCNO

Figure S7

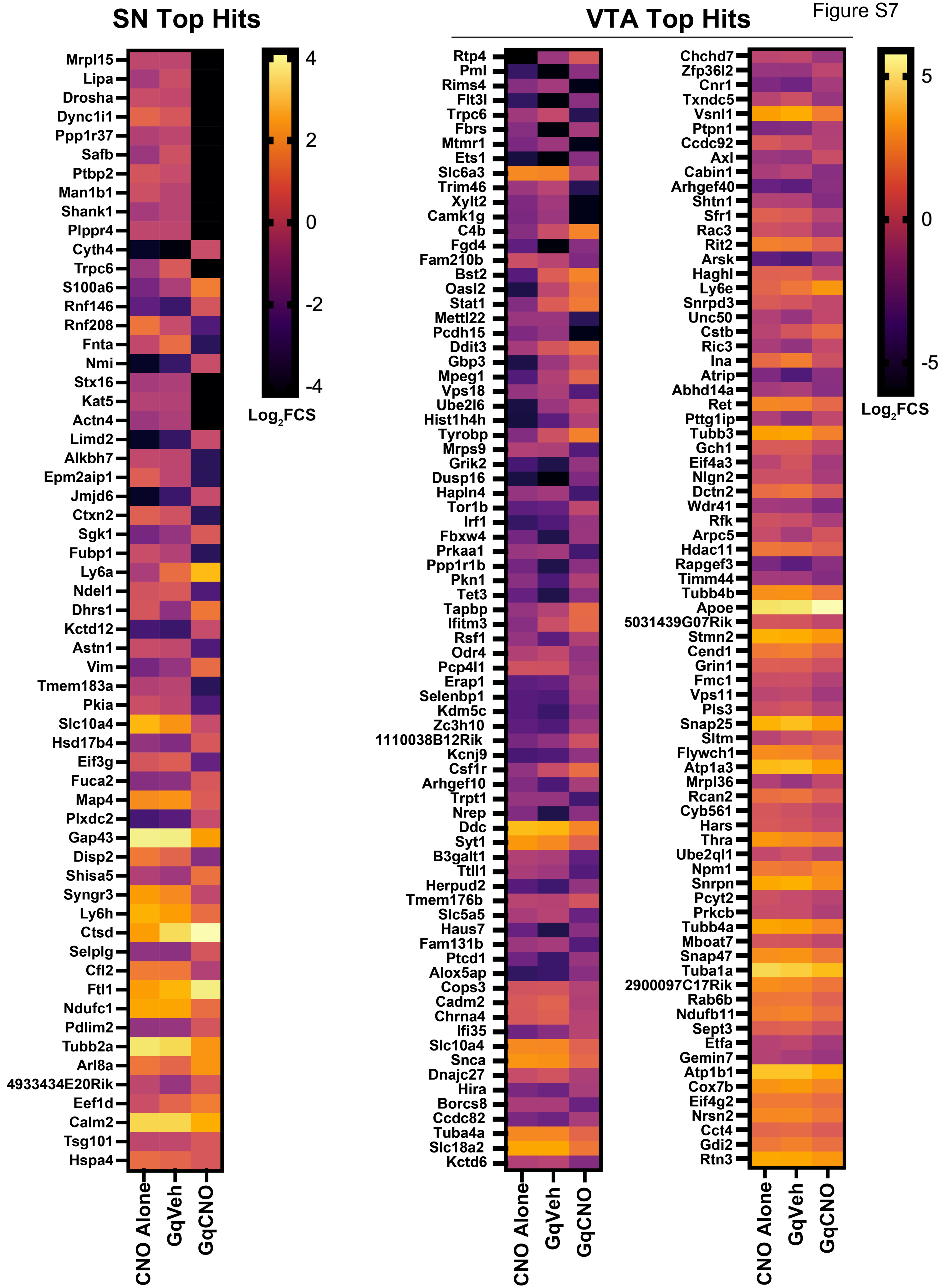
SN Top Hits
Mrpl15
Lipa
Drosha
Dync1i1
Ppp1r37
Safb
Ptbp2
Man1b1
Shank1
Plppr4
Cyth4
Trpc6
S100a6
Rnf146
Rnf208
Fnta
Nmi
Stx16
Kat5
Actn4
Limd2
Alkbh7
Epm2aip1
Jmjd6
Ctxn2
Sgk1
Fubp1
Ly6a
Ndel1
Dhrs1
Kctd12
Astn1
Vim
Tmem183a
Pkia
Slc10a4
Hsd17b4
Eif3g
Fuca2
Map4
Plxdc2
Gap43
Disp2
Shisa5
Syngr3
Ly6h
Ctsd
Selplg
Cfl2
Ftl1
Ndufc1
Pdlim2
Tubb2a
Arl8a
4933434E20Rik
Eef1d
Calm2
Tsg101
Hspa4
CNO Alone
GqVeh
GqCNO
4
2
0
-2
-4
Log2FCS
VTA Top Hits
Rtp4
Pml
Rims4
Flt3l
Trpc6
Fbrs
Mtmr1
Ets1
Slc6a3
Trim46
Xylt2
Camk1g
C4b
Fgd4
Fam210b
Bst2
Oasl2
Stat1
Mettl22
Pcdh15
Ddit3
Gbp3
Mpeg1
Vps18
Ube2l6
Hist1h4h
Tyrobp
Mrps9
Grik2
Dusp16
Hapln4
Tor1b
Irf1
Fbxw4
Prkaa1
Ppp1r1b
Pkn1
Tet3
Tapbp
Ifitm3
Rsf1
Odr4
Pcp4l1
Erap1
Selenbp1
Kdm5c
Zc3h10
1110038B12Rik
Kcnj9
Csf1r
Arhgef10
Trpt1
Nrep
Ddc
Syt1
B3galt1
Ttll1
Herpud2
Tmem176b
Slc5a5
Haus7
Fam131b
Ptcd1
Alox5ap
Cops3
Cadm2
Chrna4
Ifi35
Slc10a4
Snca
Dnajc27
Hira
Borcs8
Ccdc82
Tuba4a
Slc18a2
Kctd6
Chchd7
Zfp36l2
Cnr1
Txndc5
Vsnl1
Ptpn1
Ccdc92
Axl
Cabin1
Arhgef40
Shtn1
Sfr1
Rac3
Rit2
Arsk
Haghl
Ly6e
Snrpd3
Unc50
Cstb
Ric3
Ina
Atrip
Abhd14a
Ret
Pttg1ip
Tubb3
Gch1
Eif4a3
Nlgn2
Dctn2
Wdr41
Rfk
Arpc5
Hdac11
Rapgef3
Timm44
Tubb4b
Apoe
5031439G07Rik
Stmn2
Cend1
Grin1
Fmc1
Vps11
Pls3
Snap25
Sltm
Flywch1
Atp1a3
Mrpl36
Rcan2
Cyb561
Hars
Thra
Ube2ql1
Npm1
Snrpn
Pcyt2
Prkcb
Tubb4a
Mboat7
Snap47
Tuba1a
2900097C17Rik
Rab6b
Ndufb11
Sept3
Etfa
Gemin7
Atp1b1
Cox7b
Eif4g2
Nrsn2
Cct4
Gdi2
Rtn3
CNO Alone
GqVeh
GqCNO
CNO Alone
GqVeh
GqCNO
5
0
-5
Log2FCS

Figure S8

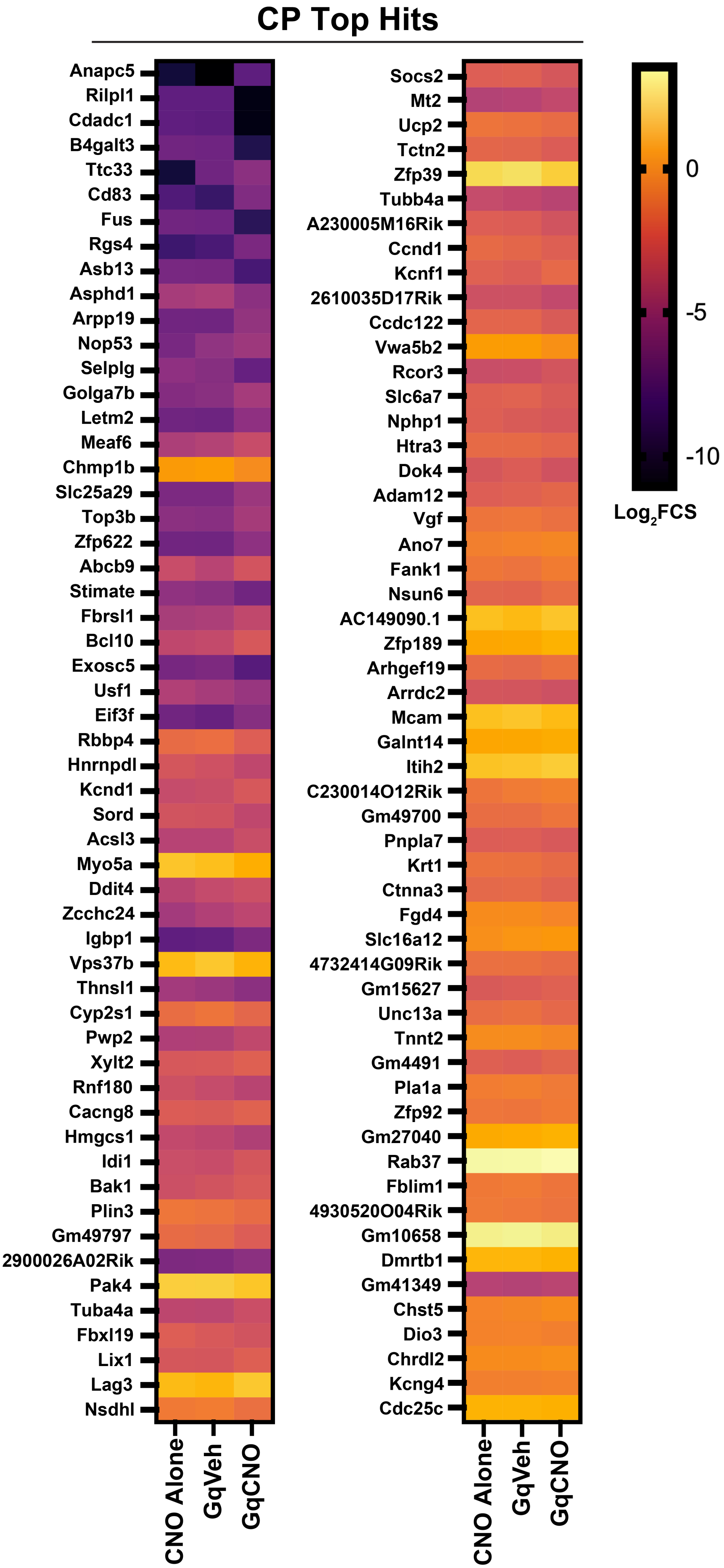

CP Top Hits
Anapc5
Rilpl1
Cdadc1
B4galt3
Ttc33
Cd83
Fus
Rgs4
Asb13
Asphd1
Arpp19
Nop53
Selplg
Golga7b
Letm2
Meaf6
Chmp1b
Slc25a29
Top3b
Zfp622
Abcb9
Stimate
Fbrsl1
Bcl10
Exosc5
Usf1
Eif3f
Rbbp4
Hnrnpdl
Kcnd1
Sord
Acsl3
Myo5a
Ddit4
Zcchc24
Igbp1
Vps37b
Thnsl1
Cyp2s1
Pwp2
Xylt2
Rnf180
Cacng8
Hmgcs1
Idi1
Bak1
Plin3
Gm49797
2900026A02Rik
Pak4
Tuba4a
Fbxl19
Lix1
Lag3
Nsdhl
CNO Alone
GqVeh
GqCNO
Socs2
Mt2
Ucp2
Tctn2
Zfp39
Tubb4a
A230005M16Rik
Ccnd1
Kcnf1
2610035D17Rik
Ccdc122
Vwa5b2
Rcor3
Slc6a7
Nphp1
Htra3
Dok4
Adam12
Vgf
Ano7
Fank1
Nsun6
AC149090.1
Zfp189
Arhgef19
Arrdc2
Mcam
Galnt14
Itih2
C230014O12Rik
Gm49700
Pnpla7
Krt1
Ctnna3
Fgd4
Slc16a12
4732414G09Rik
Gm15627
Unc13a
Tnnt2
Gm4491
Pla1a
Zfp92
Gm27040
Rab37
Fblim1
4930520O04Rik
Gm10658
Dmrtb1
Gm41349
Chst5
Dio3
Chrdl2
Kcng4
Cdc25c
CNO Alone
GqVeh
GqCNO
0
-5
-10
Log2FCS

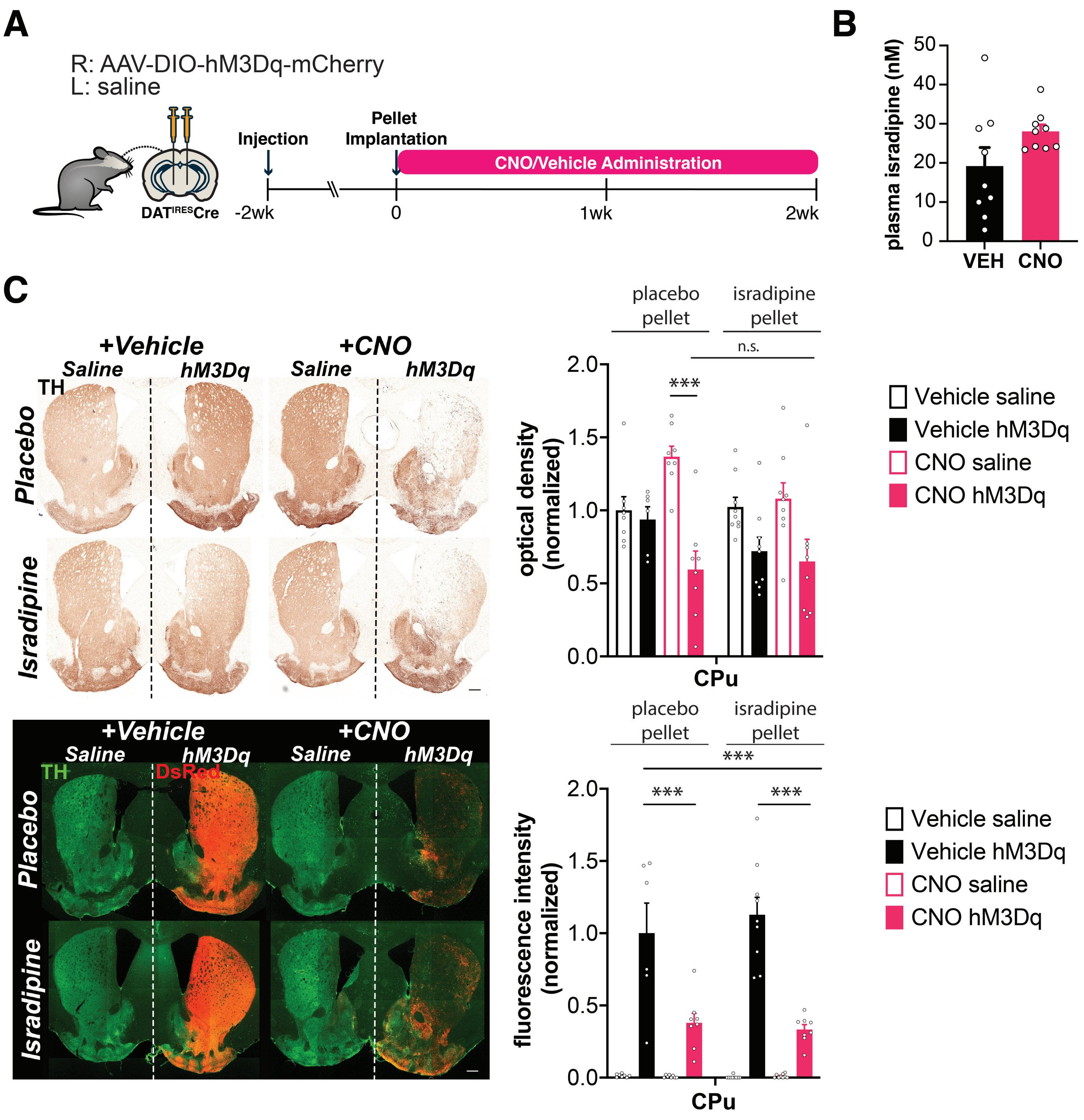
A
R: AAV-DIO-hM3Dq-mCherry
L: saline
DAT IRES Cre
Injection
Pellet Implantation
CNO/Vehicle Administration
-2wk
0
1wk
2wk
B
plasma isradipine (nM)
VEH
CNO
C
+Vehicle
+CNO
Saline
hM3Dq
TH
Placebo
Isradipine
DsRed
placebo pellet
isradipine pellet
n.s.
***
optical density (normalized)
fluorescence intensity (normalized)
CPu
Vehicle saline
Vehicle hM3Dq
CNO saline
CNO hM3Dq

Figure S10

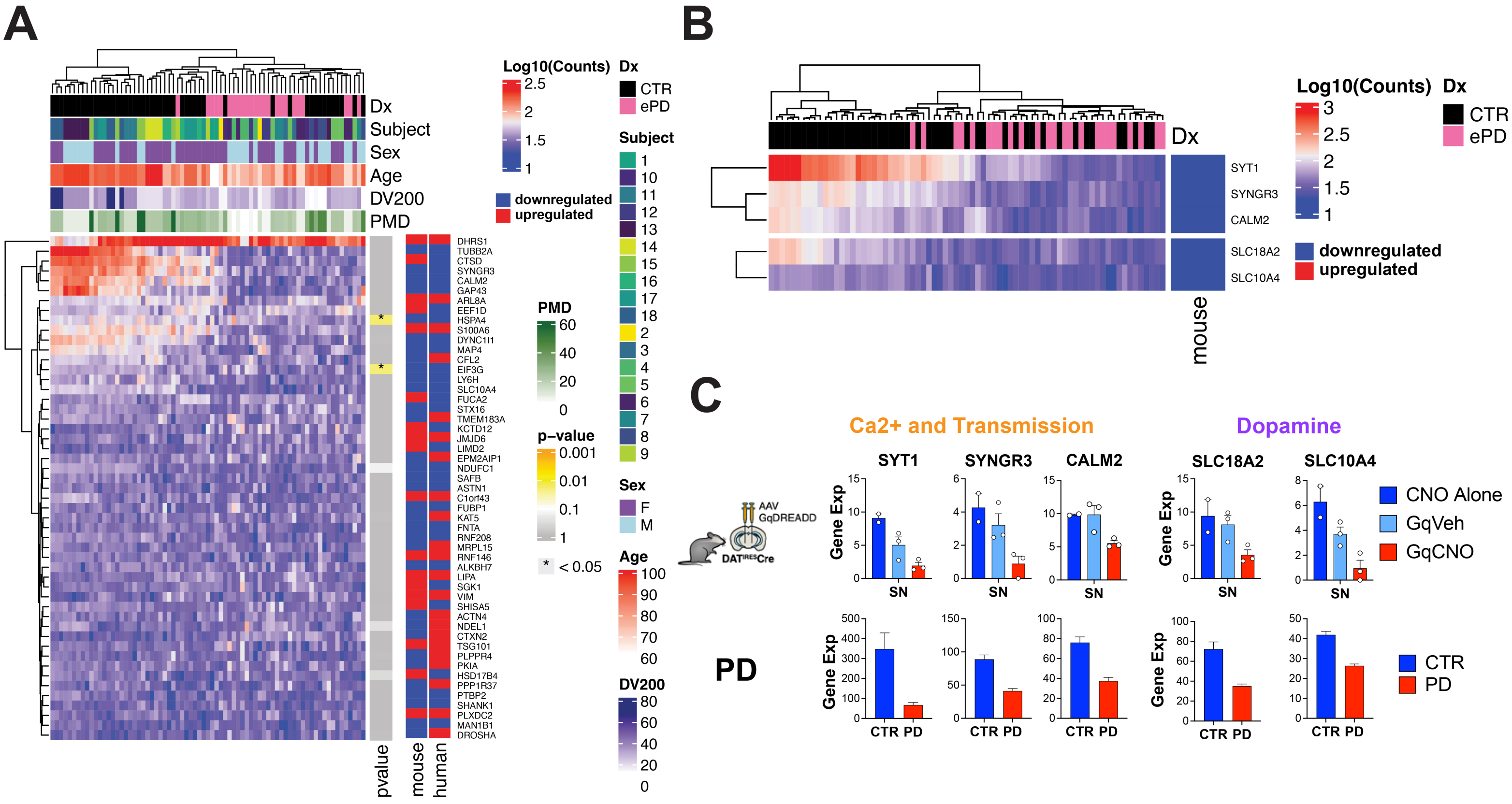

A
Dx
Subject
Sex
Age
DV200
PMD
Log10(Counts)
downregulated
upregulated
p−value
* < 0.05
CTR
ePD
F
M
pvalue
mouse
human
DHRS1
TUBB2A
CTSD
SYNGR3
CALM2
GAP43
ARL8A
EEF1D
HSPA4
S100A6
DYNC1I1
MAP4
CFL2
EIF3G
LY6H
SLC10A4
FUCA2
STX16
TMEM183A
KCTD12
JMJD6
LIMD2
EPM2AIP1
NDUFC1
SAFB
ASTN1
C1orf43
FUBP1
KAT5
FNTA
RNF208
MRPL15
RNF146
ALKBH7
LIPA
SGK1
VIM
SHISA5
ACTN4
NDEL1
CTXN2
TSG101
PLPPR4
PKIA
HSD17B4
PPP1R37
PTBP2
SHANK1
PLXDC2
MAN1B1
DROSHA
B
SYT1
SYNGR3
CALM2
SLC18A2
SLC10A4
C
Ca2+ and Transmission
Dopamine
AAV GqDREADD
DAT^IRES Cre
Gene Exp
SN
CNO Alone
GqVeh
GqCNO
PD
CTR PD

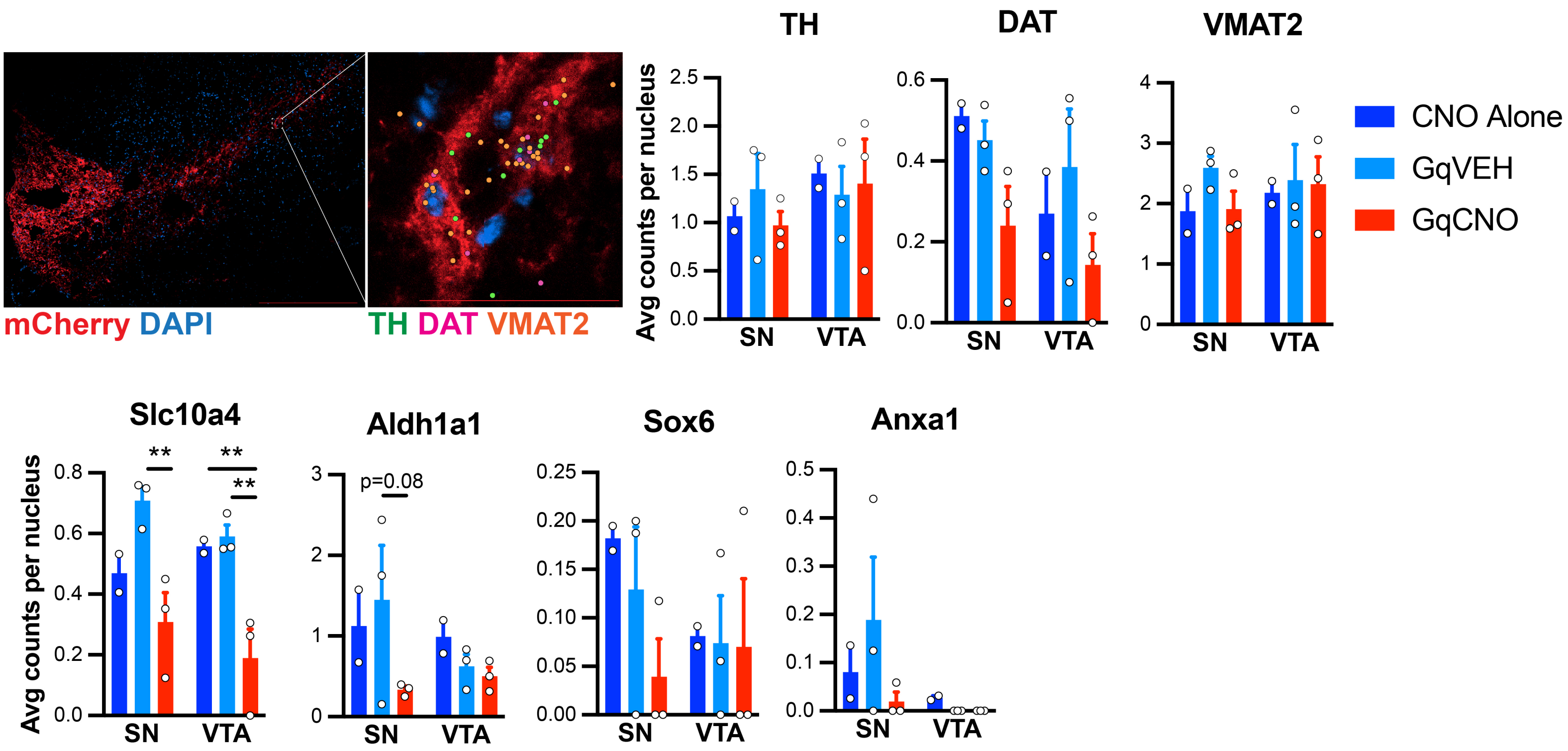
mCherry DAPI
TH DAT VMAT2
TH
DAT
VMAT2
Avg counts per nucleus
SN
VTA
CNO Alone
GqVEH
GqCNO
Slc10a4
Aldh1a1
Sox6
Anxa1
p=0.08
**

**VTA**

| Index | GO Molecular Function 2023 | P-value |
| --- | --- | --- |
| 1 | GTP Binding (GO:0005525) | 0.00002637 |
| 2 | Syntaxin Binding (GO:0019905) | 0.00005402 |
| 3 | Guanyl Ribonucleotide Binding (GO:0032561) | 0.00006568 |
| 4 | Nuclear Receptor Coactivator Activity (GO:0030374) | 0.0005911 |
| 5 | Purine Ribonucleoside Triphosphate Binding (GO:0035639) | 0.001174 |

| Index | GO Biological Process 2023 | P-value |
| --- | --- | --- |
| 1 | Chemical Synaptic Transmission (GO:0007268) | 0.00000141 |
| 2 | Synaptic Vesicle Exocytosis (GO:0016079) | 0.0000152 |
| 3 | Anterograde Trans-Synaptic Signaling (GO:0098916) | 0.00002437 |
| 4 | Response To Cytokine (GO:0034097) | 0.00005223 |
| 5 | Response To Interferon-Beta (GO:0035456) | 0.00006911 |

**SN**

| Index | GO Molecular Function 2023 | P-value |
| --- | --- | --- |
| 1 | Phosphatase Activator Activity (GO:0019211) | 0.001131 |
| 2 | Tubulin Binding (GO:0015631) | 0.002579 |
| 3 | Calcium Channel Regulator Activity (GO:0005246) | 0.005332 |
| 4 | Microtubule Binding (GO:0008017) | 0.005412 |
| 5 | Alpha-Tubulin Binding (GO:0043014) | 0.005618 |

| Index | GO Biological Process 2023 | P-value |
| --- | --- | --- |
| 1 | Positive Regulation Of Transporter Activity (GO:0032411) | 8.673E-06 |
| 2 | Establishment Of Spindle Orientation (GO:0051294) | 0.0001492 |
| 3 | Neuron Projection Morphogenesis (GO:0048812) | 0.0009083 |
| 4 | Long-Term Memory (GO:0007616) | 0.001571 |
| 5 | Vesicle Transport Along Microtubule (GO:0047496) | 0.001903 |

**CP**

| Index | GO Molecular Function 2023 | P-value |
| --- | --- | --- |
| 1 | Protein Phosphatase 2A Binding (GO:0051721) | 0.007033 |
| 2 | Histone Deacetylase Binding (GO:0042826) | 0.01734 |
| 3 | Calcium Channel Regulator Activity (GO:0005246) | 0.01761 |
| 4 | Protein Phosphatase Binding (GO:0019903) | 0.02563 |
| 5 | arachidonate-CoA Ligase Activity (GO:0047676) | 0.0272 |

| Index | GO Biological Process 2023 | P-value |
| --- | --- | --- |
| 1 | DNA Deamination (GO:0045006) | 0.001908 |
| 2 | Positive Regulation Of Mitotic Cell Cycle Phase Transition (GO:1901992) | 0.005059 |
| 3 | Positive Regulation Of G2/M Transition Of Mitotic Cell Cycle (GO:0010971) | 0.00588 |
| 4 | Positive Regulation Of Cell Cycle G2/M Phase Transition (GO:1902751) | 0.007645 |
| 5 | Secondary Alcohol Biosynthetic Process (GO:1902653) | 0.007645 |

**Supplementary Table 2** - Demography of the human post-mortem cohort assayed by GeoMx.

| Group | Gender (M/F) | Age at death (years) | Post-mortem delay (hours) | DV200 |
|---|---|---|---|---|
| Aged Healthy Controls | 3/7 | 92.0 (5.75) | 25 (8.25) | 28.9 (16.4) |
| Early PD | 5/3 | 73.5 (12)** | 17.5 (13.5)* | 28.3 (9.10) |

Values are presented as median (IQR). The comparison of Age at death, Post-mortem delay and DV200 between groups was made using the Welch Two Sample t-test. The comparison of gender between groups was made using chi-square test. * $P < 0.05$, ** $P < 0.001$.
